# On the use of variational autoencoders for biomedical data integration

**DOI:** 10.1101/2025.08.18.670835

**Authors:** Marc Pielies Avellí, Ricardo Hernández Medina, Henry Webel, Simon Rasmussen

## Abstract

Variational Autoencoders (VAEs) are a widely used framework to integrate diverse biomedical data modalities, create representations that capture the underlying structure of the datasets, and obtain insights about the relations between variables. Here we describe how this is achieved from an empirical point of view in our previously developed VAE-based framework MOVE, providing an intuitive perspective on the inner workings of multimodal VAEs in biomedical contexts. We explore how the models’ emerging dynamics shape their performance and how *in silico* perturbations can be leveraged to identify potential associations between variables. To do that, we extend our framework to handle perturbations of continuous variables, introduce a new approach to better capture associations between them, and create synthetic datasets to benchmark the proposed methods against well-defined ground truth associations. We finally showcase our findings in real biomedical scenarios, namely a multimodal dataset of inflammatory bowel disease and a dataset containing genetic knockdowns in K562 and RPE1 cells.

## Background

Life sciences research often yields high-dimensional, noisy, and multi-modal datasets from a limited number of biological samples [1,2]. Integrating these complementary data types into meaningful and interpretable representations is hence of utmost importance when trying to understand complex biological systems [3].

Deep neural networks, particularly Variational Autoencoders (VAEs), are well-suited for this data integration task [4]. A VAE consists of an encoder that compresses high-dimensional inputs into a lower-dimensional latent representation, and a decoder that reconstructs the original data from this latent space. The model is trained to minimize the reconstruction error while organizing the samples in latent space according to a prior distribution. VAEs have been applied to diverse biomedical challenges, including patient stratification [5], single-cell data embedding [6,7], and metagenome binning [8,9]. Our previously developed multi-omics variational autoencoder (MOVE) integrates continuous and categorical biomedical data, as demonstrated on a cohort of type 2 diabetes (T2D) patients [10].

VAEs learn the intricate relationships between input variables, making them a powerful framework to perform *in silico* perturbations. For example, the compositional perturbational autoencoder (CPA) modeled the effects of chemical perturbations by combining the latent representations of cells, covariates, and drugs [11]. Conversely, MOVE simulated perturbations by directly modifying input data, such as changing a patient’s medication status, and measuring the resulting changes in the reconstructed outputs [10]. To identify significant associations, Allesøe et al. developed two statistical methods: a t-test on reconstructed feature values and a Bayesian-inspired test to detect significant shifts in feature distributions after the perturbations [10]. These methods were originally developed to handle perturbations of categorical variables, but many potential applications required their extension to continuous data, e.g. modeling the effects of an increased concentration of a metabolite or protein in a sample.

In this manuscript, we extend the existing MOVE framework to facilitate the analysis of continuous variable perturbations and introduce the Kolmogorov-Smirnov (KS) distance as a novel metric to capture the magnitude and consistency of reconstruction shifts. We provide new methodological insights, moving beyond the original MOVE setup to offer a mechanistic perspective on how these models behave internally, bridging the gap between high-level performance and latent space dynamics (**Figure 1**). We showcase our findings in synthetic datasets, evaluate the data integration ability of the model on a multi-omics study of Inflammatory Bowel Disease (IBD) [12], and benchmark the model’s predictive performance using Perturb-seq data on K562 and RPE1 cells from Replogle et al. [13]. These complementary analyses provide a grounded understanding of VAEs’ inner workings, their power and limitations when applied to diverse biomedical contexts and tasks.

**Figure 1.**
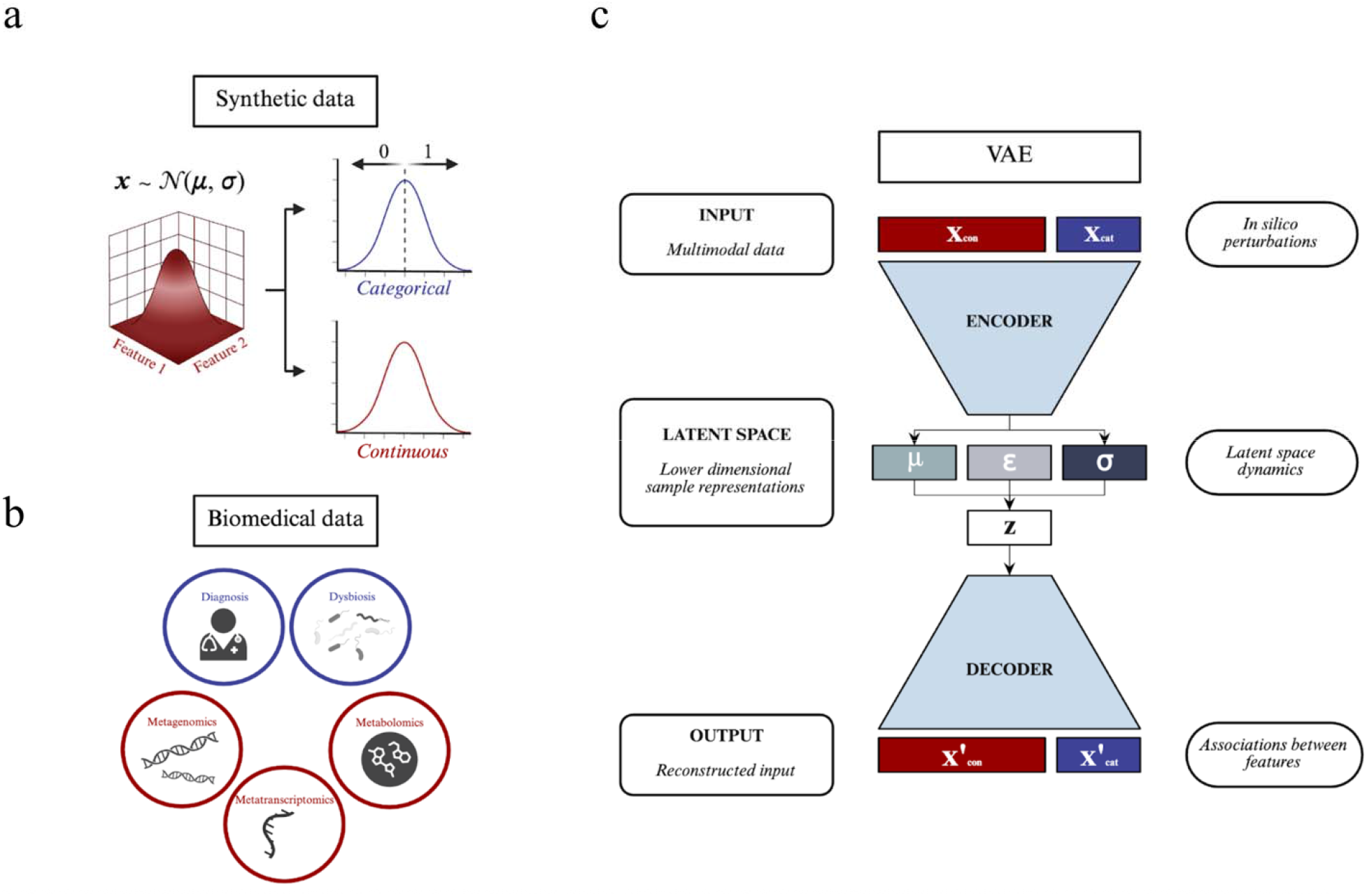
Graphical summary. **a)** Synthetic data generation. Samples are drawn from a multivariate gaussian distribution, where the degree of association between variables is defined to be their correlation. **b)** Biomedica multimodal data can contain categorical features (e.g. the diagnosis of a disease), described by classes, and continuous features (e.g. metabolite or RNA counts in a sample), described by continuous values. **c)** Variational autoencoders (VAEs) can compress high-dimensional data into lower dimensional representations. The trained models can then be used for downstream tasks such as modeling the impact of *in silico* perturbations, identifying key features and associations between them.

## Results

### The properties of the dataset shape the behavior of the VAE

Assessing a VAE’s ability to capture biological dynamics is challenging without access to ground truths. To address this, we created synthetic datasets that emulated multimodal readouts from a set of samples with measured continuous and categorical properties. Continuous features represented, for instance, metabolite concentrations, while binary features simulated variables like the presence of a metagenomic species or a patient’s disease status. The protocol generated low-sparsity datasets where the strength of associations between features was defined by the correlation coefficient (**Methods**).

A simple, fully connected VAE was trained on these synthetic datasets (**Figure 2a**). While spurious correlations emerged with few samples (I), they disappeared when more samples were used (II) (**Supplementary Figure 1**). Under a low correlation regime (II), the models learned a limited number of input features, constrained by the network’s bottleneck, as independent inputs could not be easily compressed (**Figure 2bII**). This aligns with Elhage et al. [14], who noted that separate latent dimensions are needed for each independent input component (**Supplementary Figure 2**).

**Figure 2.**
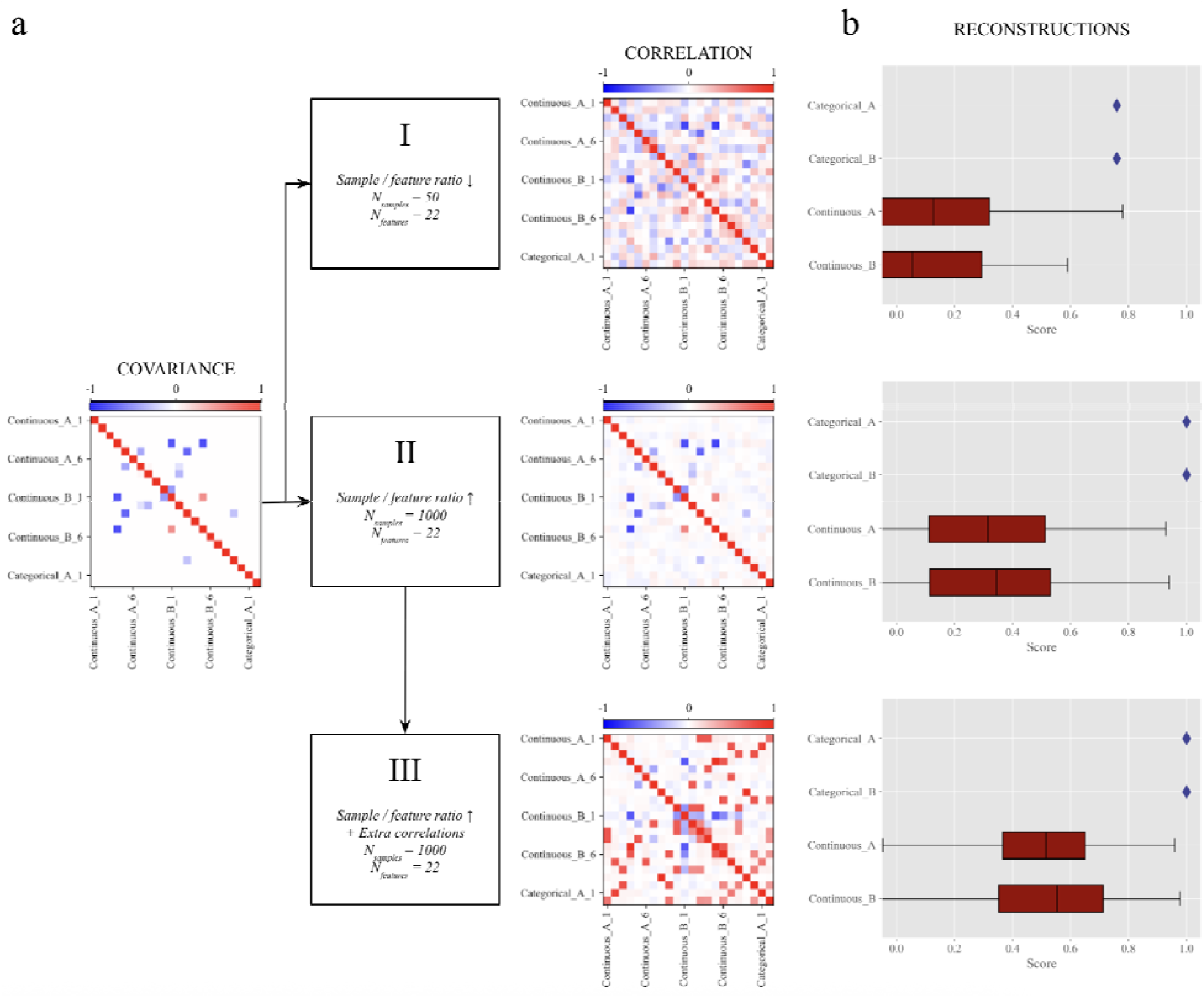
The behavior of the VAE depends on the nature of the dataset. The same VAE architecture (N_hidden_ = 20, N_latent_ = 5 and small regularization (β = 0.001)) was trained to compress and reconstruct three datasets. **a)** Ground truth correlations between features under a sparse covariance regime for I) low sample to feature ratio, II) high sample to feature ratio and III) high sample to feature ratio with added explicit correlations between variables. **b)** Normalized reconstruction accuracies for categorical and continuous features for all samples in each set. A categorical variable was properly reconstructed if the class with the highest probability corresponded to the class given by the input. Accuracy in continuous feature reconstructions was assessed by comparing the reconstructed array with the input array using cosine similarity for each individual. Reconstruction accuracies increased when adding correlations between features.

However, in most biological systems, variables are entangled to some degree and therefore there is some redundancy (e.g., gene co-expression patterns). This means the data lies on a lower-dimensional manifold, which allows a VAE to compress and reconstruct it more effectively (**Figure 2bIII**). Multicollinearity, common in biomedical data where features outnumber observations, also contributes to lowering the data’s dimensionality, as some variables can be written as linear combinations of others [1].

In summary, a VAE’s ability to compress and reconstruct data highly depends on the dataset’s properties, including the information shared between variables, sparsity, and the feature-to-sample ratio. It is therefore crucial to assess whether the model has learned a feature before using it for downstream tasks, such as feature importance or perturbation analyses.

### MOVE arranges samples in latent space to ease their reconstructions

Variational Autoencoders (VAEs) were originally designed to understand the underlying generative process of the data by maximizing the evidence lower bound (ELBO). This objective function implicitly encoded two tasks: maximizing the reconstruction accuracy for both categorical and continuous variables, while imposing a certain regularization on the latent space (**Methods**). All subsequent results, from latent space analyses to the identification of associations between variables stemmed from this training objective.

To facilitate the reconstruction of the samples, the model’s encoder tried to separate them into different clusters in the latent space based on shared categories or labels (**Figure 3a**). This eased the task for the decoder, which acted as a multi-task classifier. In a similar fashion, the encoder ordered samples along value gradients of the learned continuous variables (**Figure 3b-d**). On the other hand, variables that the model could not encode did not display any trend. Samples displaying similar values for these features were scattered across the latent space. Having samples with similar values scattered around would complicate the task for our rather simple decoder, resulting in poor feature reconstructions (**Figures 3e-f**). Thus, measuring a high feature reconstruction could be used as a proxy for a meaningful latent representation of said feature.

**Figure 3.**
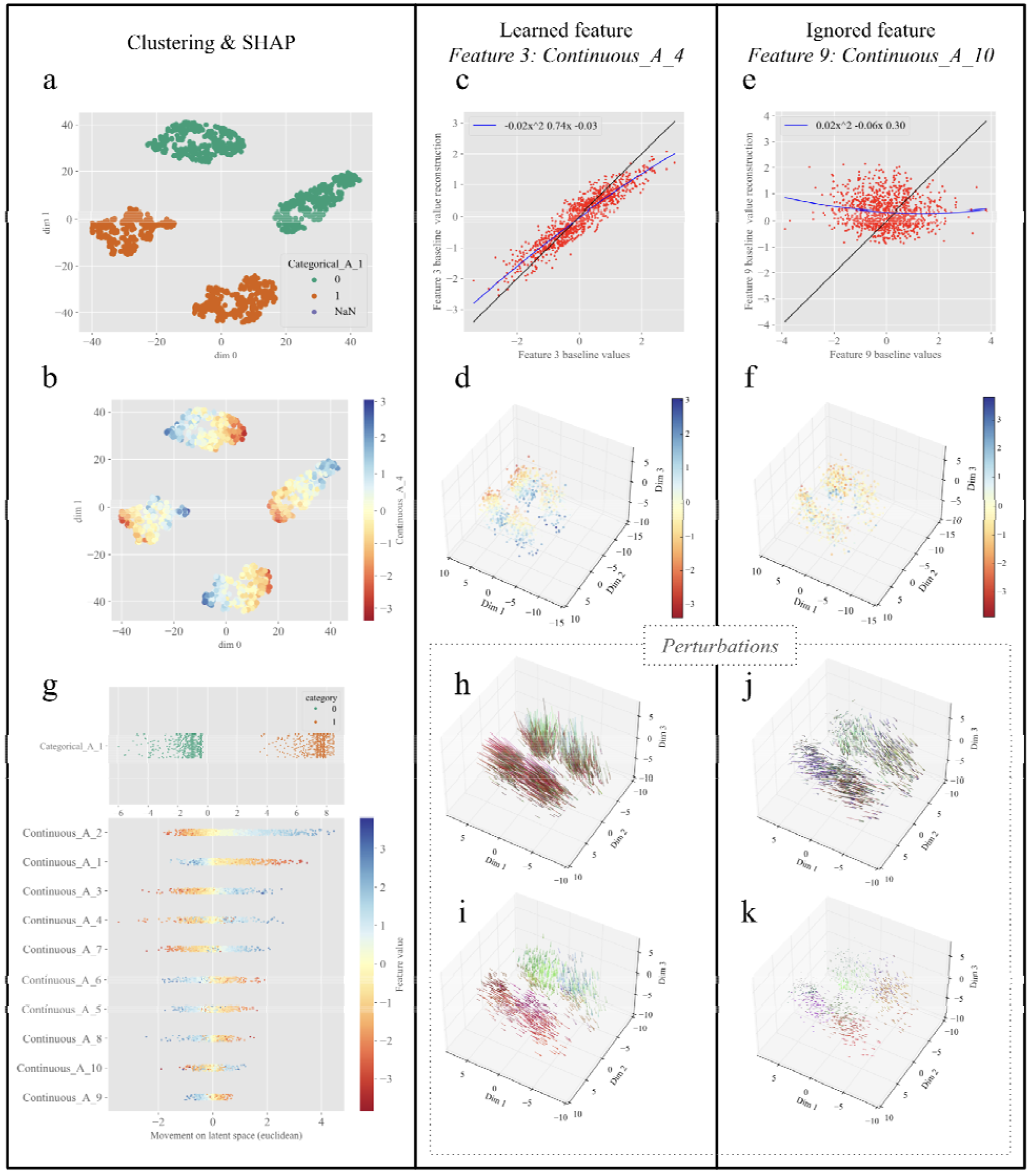
The distribution of samples in latent space shapes all downstream analyses. Samples were arranged in a latent space **a)** in clusters according to the learned categorical features and **b)** following value gradients of the learned continuous features. **c)** Reconstructed values vs. input values for a learned input variable (Feature 3, Continuous_A_4). **d)** Untransformed visualization of a 3D latent space (N_latent_ = 3) color coded by the feature values of a learned input variable (Feature 3, Continuous_A_4). **e)** Reconstructed values vs. input values for a feature the model ignores (Feature 9, Continuous_A_10). **f)** Latent space color coded by the feature values of an ignored input variable (Feature 9, Continuous_A_10). **g)** SHAP plots for categorical variables (top) and continuous (bottom) ranking features according to their importance to the model. Each dot represents a sample, color coded by its value of the feature of interest. Its x-axis location is the Euclidean distance travelled by the sample in latent space when setting the value to zero. **h-i)** Movement of the latent space representation of the samples, with arrowheads at the location of the sample after the perturbation, tails at the location of origin. Movement when setting the value of a learned perturbed feature to **h)** the maximum found in the cohort or **i)** adding a small quantity to the original value. **j-k)** Movement of the latent space representation of the samples when perturbing an ignored feature by **j)** setting its value to the maximum or **k)** adding a small quantity.

These categorical-driven and continuous-driven ordering mechanisms gave rise to a spectrum of ordering regimes, determined by the importance assigned to each data modality. At one extreme, samples were highly clustered to ensure perfect categorical reconstructions. This clustering, however, produced a sparsely populated latent space (**Figure 3a, b**). On the other end, categorical variables would be ignored to foster the reconstruction of continuous datasets, inducing a more homogeneous latent space (**Supplementary Figure 3**). The relative importance of each dataset thus constituted a key hyperparameter that determined the model’s behavior. Finally, a Gaussian prior acted as an effective force pushing samples towards the center of the coordinate system.

In summary, the encoder and decoder networks worked together to maximize the model’s reconstruction accuracy. The encoder distributed samples into clusters according to categorical variables and along the value gradients of learned continuous variables. To promote a densely populated latent space and prevent excessive separation of samples, the contribution of continuous variables could be upweighted and regularization could be enforced. This information was exploited in the following sections.

### SHAP analysis reflects the organization of samples in latent space

We proceeded to identify which input features were relevant to characterize the system from the model’s perspective using a SHAP-inspired analysis (**Figure 3g**). The intuition was that if a feature was important to the model, its removal would highly affect a sample’s latent representation. To quantify this, we set each feature’s value to zero across all samples, one by one, and measured the resulting movement of samples in the latent space. An importance ranking was then derived from the cumulative impact each feature had on the samples’ latent space locations (**Methods**).

The arrangement of samples in the latent space presented in the previous section deeply shaped the results obtained from the feature importance analysis. The highest importance was assigned to features that, when removed, induced greater and more homogeneous changes in sample locations (**Figure 3g**). These were the features the model had learned, either by spreading samples along value gradients (e.g., Continuous_A_2 or A_4) or by grouping them into class clusters (e.g., Categorical_A and B). Samples with similar values for these learned features moved similarly when perturbed (**Figure 3g, top, Methods**). Conversely, removing features that were not important to the model resulted in samples remaining close to their original positions (**Figure 3g, bottom**). This was the case for unlearned features (e.g., Continuous_A_10) but also those that were learned but varied over short latent distances.

In conclusion, a SHAP-based analysis provided insights into the samples’ organization in latent space and offered a first hint at which variables the model had learned.

### Context-preserving perturbations provide robust changes in the latent space

*In silico* perturbations constitute a promising technique to study the relations between variables captured by the models. A desirable property of such perturbations would be to keep the directionality of the change across samples, which was not the case for the SHAP analyses.

Our initial perturbation method involved setting a feature’s value across all samples to the cohort’s minimum or maximum. However, this approach did not affect all samples uniformly: setting a feature to the cohort maximum, for example, was highly disruptive for samples with low initial values, causing a large jump in their latent space representation (**Figure 3h**). On the other hand, samples with values already close to the maximum were less affected. Arbitrarily large perturbations risked moving samples into unexplored latent regions, creating out-of-distribution inputs. Such extreme inputs may additionally disrupt the learned data structure. Because the model was trained only on plausible, unperturbed data, its reconstructions of these displaced samples could lack biological meaning.

The non-uniform effects and out-of-distribution risks of value-substitution perturbations led us to propose an alternative scheme that preserved the feature’s original context. To achieve this, we added or subtracted a fixed quantity, namely the standard deviation (i.e. 1 after normalization) from the input feature values. Samples moved in a smoother manner than in the value substitution scenario, staying around the sampled region of the latent space (**Figure 3i**). As in the SHAP-based protocol, the movement of samples when perturbing unlearned features was smaller under both perturbation schemes (**Figures 3j,k**). Such movements followed the direction defined by the gradient of feature values in the local latent neighborhood. Finally, not only the directionality but also the magnitude of these perturbations played a key role in the latent displacements of the samples and therefore in their reconstructions.

The choice between these perturbation methods should be dictated by the biological context and the risk of out-of-distribution effects. Substituting a feature with the cohort’s minimum or maximum value is suitable e.g. for simulating “knock-out” scenarios. Yet, it risks pushing samples into unexplored latent regions where the decoder may produce biologically implausible reconstructions. Conversely, adding or subtracting one standard deviation (±1*SD*) serves as a context-preserving scheme that models physiological variations while reducing the risk of samples leaving the learned manifold, avoiding extreme latent displacements.

### Measuring perturbation effects on the reconstructed samples using statistical tests

Having established that context-preserving perturbations produce consistent, directional movements in latent space, we next studied how such latent shifts affected sample reconstructions and how the resulting output changes could reveal associations between variables.

Perturbing learned variables induced systematic changes in the reconstructions of their associated variables, which could be used to identify their relationships. For instance, a positive input perturbation moved samples to latent regions with higher feature values, increasing the reconstructed values of positively correlated variables (**Figure 3h-i, Supplementary Figure 4**). In our original MOVE framework [10], we proposed two methods to quantify these changes, but they had notable limitations. A t-test comparing baseline and perturbed reconstructions produced a high false-positive rate (**Figure 4a, Supplementary Figure 5**). In contrast, a Bayes-inspired method achieved better precision but did not quantify the strength of an association (i.e. its effect size). Instead, it scored the likelihood of replication across samples. Associations for which all samples replicated the direction of change received the maximum attainable score, making them difficult to prioritize (**Figure 4b top, Supplementary Figure 5**).

**Figure 4.**
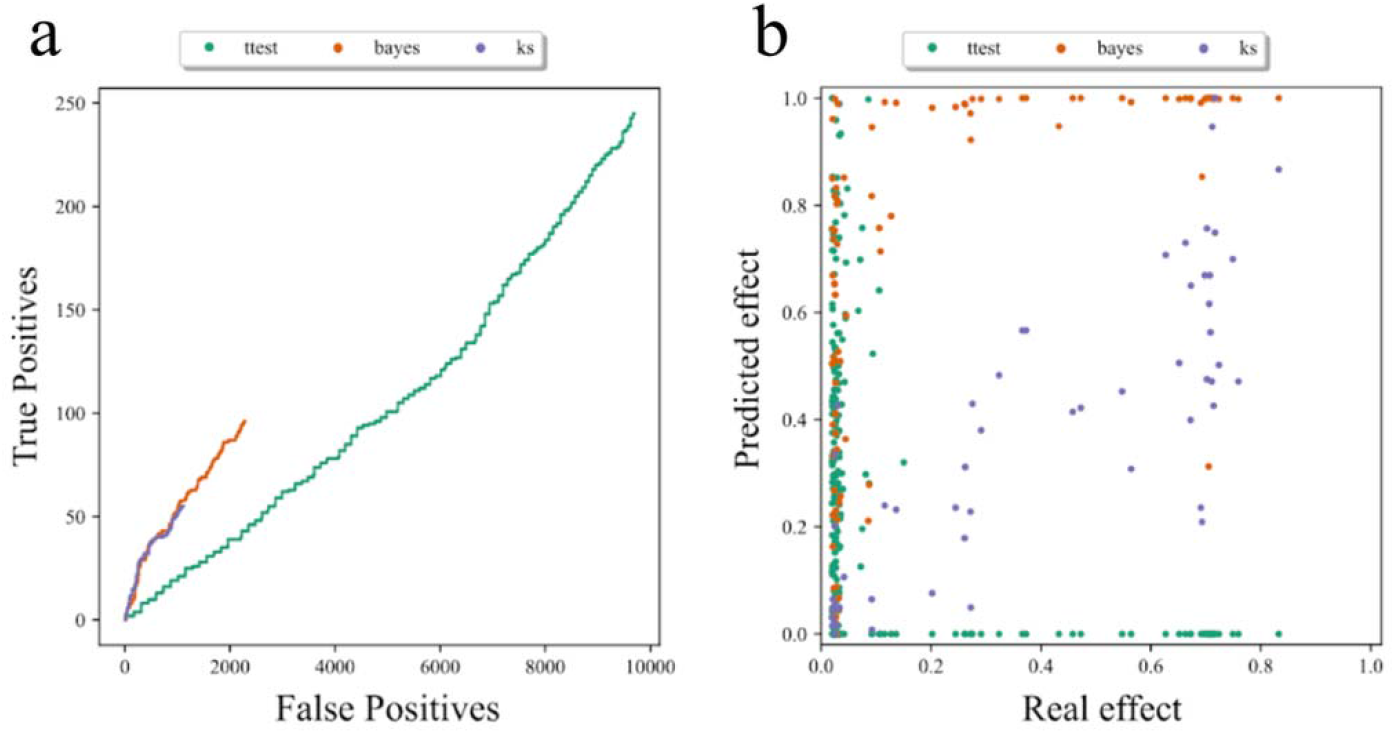
Comparison of methods to identify associations between variables. Dataset with N_samples_ = 5000 N_features_ = 150 with added correlations (III). **a)** True positive vs false positive association discovery when reading sequentially the lists of found associations, ranked by effect size, for the different methods. **b)** Predicted vs. Real effect for the true positive associations found by the three methods. The effect strength, i.e. the real effect, was set to be the correlation coefficient between the two associated features. Predicted Bayes and KS scores were linearly transformed between 0 and 1. T-test predicted effects were calculated as -log_10_(p-value) and were also linearly scaled after the transformation. Note a saturation of highly important associations in Bayes and that KS scores for correctly predicted associations (predicted effect) correlate with association strength (real effect).

**Figure 5.**
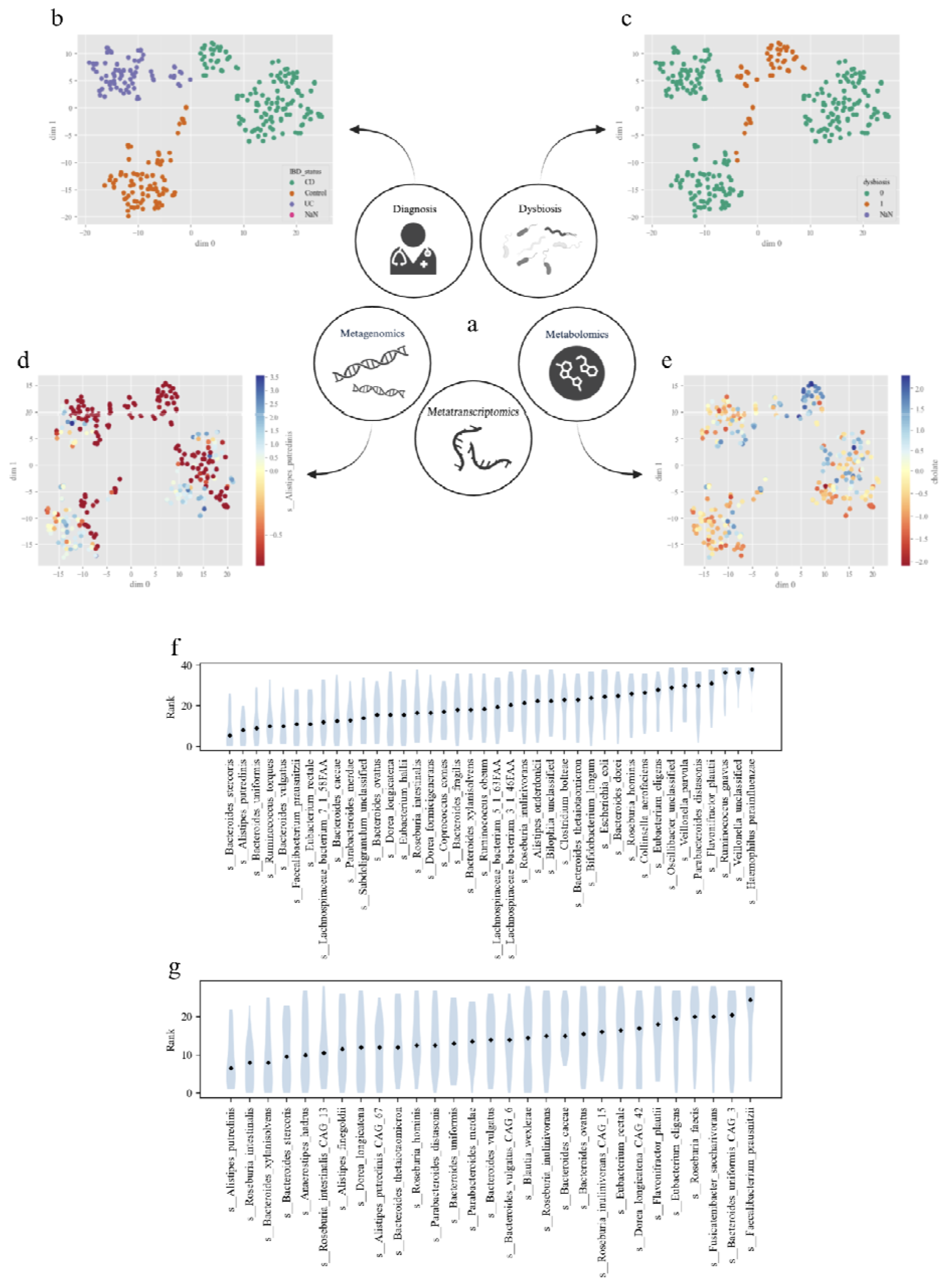
Multi-omics data integration in the context of Inflammatory Bowel Diseases. **a)** Overview of the data modalities included in our analyses. The datasets contained 293 samples and the following number of features: # Metabolomics = 592, # Metagenomics = 40, # Metatranscriptomics = 29, # IBD status = 1 (3 categories → 3 input nodes), # Dysbiosis = 1 (2 categories → 2 input nodes). **b-e)** t-SNE of the latent embeddings of a MOVE model with N_latent_ = 16, N_hidden_ = 64 and β = 0.0001, color coded by **b)** Disease status, **c)** Dysbiosis status, **d)** *Alistipes putredinis* metagenomics log-norm counts, example of continuous feature completely depleted in dysbiotic samples, **e)** log-norm cholate levels. **f, g)** SHAP ranking distribution of each feature across the following hyperparameter combinations: N_latent_ = {16, 32}, N_hidden_ = {64, 256, 512} and β = {0.01, 0.0001}. **f)** Metagenomics. **g)** Metatranscriptomics. Features on the left were consistently prioritized by the models, regardless of specific hyperparameter choices.

### Kolmogorov-Smirnov distances can be used to identify associations

To quantify the effect size of an association, we developed a third method using the Kolmogorov-Smirnov (KS) distance. This distance was the maximum vertical difference between the cumulative distribution functions (CDFs) of a feature’s reconstructed values before and after a perturbation. The resulting KS score captured both the consistency of the change across samples but also the magnitude of such change. While the KS method behaved similarly to the Bayesian method at low recall, its theoretical significance cutoff was stricter (∼ N _samples_^-½^), leading to more stringent filtering and fewer detected associations (KS_TP_ = 55, Bayes_TP_ = 96) (**Figure 4a, Methods**). The new approach however improved the ranking of important associations by more rarely assigning the maximum possible score (Max Score = 1) (**Figure 4b Supplementary Figure 5d-e**).

In summary, by quantifying systematic changes in the reconstructions after input perturbations using various statistical tests, we were able to identify putative associations between features. It is important to emphasize that different statistical approaches serve different analytical needs. The Bayesian-inspired method is highly sensitive and can identify very small shifts as significant provided they are consistent across the cohort, which may lead to a higher number of detected associations. In contrast, the KS distance requires a larger distributional shift to reach significance. Because its threshold is strictly dependent on sample size 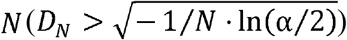, it acts as a more conservative filter that prioritizes associations with larger effect sizes.

### Multi-modal analyses of inflammatory bowel disease data

To illustrate the implications of the observed latent space dynamics and the power of our methods in a real biomedical setting, we analyzed a dataset describing the gut microbial ecosystem in inflammatory bowel diseases (IBD) [12] (**Figure 5**). IBD, which includes Crohn’s disease (CD) and ulcerative colitis (UC), are conditions characterized by inflammation of the gastrointestinal tract and are widely associated with an imbalanced gut microbiome. The multiomics dataset from Lloyd-Price et al. contained metagenomics, metatranscriptomics, and metabolomics data, along with the patients’ diagnosis and dysbiosis status [12] (**Figure 5a**).

As seen previously, MOVE clustered samples according to the categorical variables, namely disease and dysbiosis status for this cohort (**Figure 5b-c**). The encoder created simple boundaries between classes, which enabled the decoder to perfectly reconstruct these categorical features and minimize their contribution to the loss (**Supplementary Figures 6, 7**). The decoder could still reconstruct the categorical labels of held-out samples in a cross-validation setting, even when these flags were not provided as inputs to the model (**Supplementary Figures 8-11, Supplementary Table 1**). Therefore, the rest of the omics variables contained enough information about dysbiosis and disease status to enable the classification by the decoder. A baseline principal component analysis (PCA=16 components) of the joint continuous profiles (metagenomics, metatranscriptomics and metabolomics) further confirmed these results: mapping the PCA representations of the samples to the corresponding dysbiosis and disease status using a logistic regression yielded a similar classification performance (**Supplementary Figure 12, Supplementary Table 1**). Of note, the reconstructability of categorical features based on continuous features is highly dataset-dependent and was not observed in the original MOVE study [10].

The observed latent space organization both shaped and confounded subsequent analyses. To correctly reconstruct categories, the model pushed samples apart (Δ *d*_*Dysb*.*clust*.*centroid*_ ∼ 12 *l*.*u*., latent units or euclidean distance between dysbiosis cluster centroids in 32 latent space dimensions). In doing so, it also widened latent space distances for continuous features entangled with those categorical variables. Perturbing these continuous features therefore resulted in larger sample movements and higher SHAP rankings. This was observed for bacteria depleted in dysbiotic samples, like *Alistipes putredinis*, where perturbations induced larger movements (0.54 l.u.) compared to less impactful bacteria like *Veillonella* (0.28 l.u.) (**Figure 5d**,**f-g, Supplementary Table 2**). Once the categorical classification was achieved, the model continued to arrange samples based on the gradients of continuous variables, such as the metabolite cholate (**Figure 5e, Supplementary Figure 7**).

In summary, we used MOVE to integrate multi-omics data from an IBD study. The model combined information from metagenomics, metatranscriptomics, metabolomics, and patient status into a meaningful, lower-dimensional representation that facilitated data interpretation.

### The predicted associations recapitulate known biology

Our perturbational methods recapitulated the known biological associations from the original Lloyd-Price et al. analysis. For instance, perturbing the dysbiotic status of non-dysbiotic samples led to increases in the primary bile acid cholate (Bayes Score BS = 2.37) and decreases in the secondary bile acids litocholate (BS = -2.17) and deoxycholate (BS = -2.06) in the reconstructed samples (**Figure 5e, Supplementary Table 3**). These results were robust to different model initializations and training data partitions (**Supplementary Table 3**). Our perturbational approach also confirmed the previously reported associations with carnitines (BS_C14:2 Carnitine_ = 2.33, BS_C12:1 Carnitine_ = 2.21, BS_C10 Carnitine_= 2.10, etc.) (**Supplementary Tables 3**,**4**). Furthermore, carnitines were found to be associated with *Roseburia hominis* levels when perturbing both its metatranscriptomics (e.g. BS_C16 Carnitine_=-2.85, BS_C9 Carnitine_=-2.85) and the metagenomics (e.g. BS_C18:1-OH carnitine_=-4.98, BS_C16-OH carnitine_=-4.28) profiles. The strong coupling between metagenomics and metatranscriptomics for the same species promoted many cross-modality associations. Such coupling was especially pronounced for *Roseburia intestinalis, Alistipes putredinis*, and *Parabacteroides merdae*, whose metagenomic and metatranscriptomic profiles were highly correlated (**Supplementary Tables 3**,**4, Supplementary Figure 13**).

### Linking perturbation effects with association significance in the IBDMDB dataset

Despite the biological soundness of the identified associations, the perturbed multi-modal reconstructions deviated only slightly from their baselines, with KS distances around 0.08 (**Supplementary Table 4**). For our sample size (N=293, α=0.05), the KS significance threshold was approximately 0.11, meaning most observed shifts fell below it. Multiple testing corrections would further constrain detection (**Supplementary Figure 14, Methods**). In contrast, the more relaxed definition of the Bayes-inspired method considered these same small shifts to be significant (**Supplementary Tables 3**-**5, Supplementary Figure 15**).

Similar principles applied to categorical perturbations. For instance, perturbing the dysbiosis status moved samples in the latent space 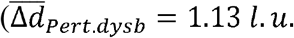, mean displacement across samples), but this was insufficient to cross the large gap between clusters (Δ*d*_*Dysb*.*clust*.*centroid*_ ∼ 12*l*.*u*.) (**Supplementary Table 2**). Consequently, the reconstructed profiles did not reflect the perturbed status and instead reverted to the original “non-dysbiotic” category (**Supplementary Table 5**). These findings highlight a limitation of the current models and urge caution when interpreting the predicted impact of *in silico* perturbations as the expected biological outcomes of such interventions.

In summary, different definitions of an association yielded complementary interpretations. The significance of an association was heavily influenced by factors like sample size, perturbation magnitude, and latent space density. Thus, we would suggest using Bayesian-inspired scores when looking for consistent changes across samples independently of their magnitude. Conversely, we would use KS-based scores to prioritize trends based on effect sizes, which should be confirmed in larger and complementary cohorts. Regardless of the perturbation size, the proposed KS method successfully ranked the degree of association between variables, allowing for their prioritization in downstream analyses.

### Performing *in silico* perturbations on VAEs in the context of the virtual cell

Another biomedical problem that has lately drawn a lot of interest is the creation of a virtual cell. A reliable digital twin of the cell could be used to infer the outcomes of unseen chemical and genetic perturbations, lowering the time and economical resources required to perform the same experiments in the lab [15,16]. We therefore wanted to explore the use of *in silico* perturbations in the input of our VAEs to predict the impact of a genetic knock-down (KD) on the cell’s transcriptome, a standard evaluation task in the field.

To that end, we trained MOVE to compress and reconstruct Replogle’s perturb-seq experiments on scRNA-seq data from RPE1 and K562 cell lines across multiple regularization regimes (β ∈ {1e-4,1e-2,1}) [13,17]. Once the model had learned the relations between gene expression levels across pseudobulk profiles for the different knock-down conditions, we used the trained model to predict the impact of lowering (minus_std) or removing (minimum) the expression of held-out knocked-down genes on the overall transcriptome (**Methods**). We followed a 4-fold validation scheme mimicking the data partitioning percentages in Ahlmann-Eltze et al. [18]. We then validated MOVE’s performance against the presented models using the standard L2 error and Pearson evaluations [11,17–23] (**Methods**). Input perturbations were applied gene by gene to control pseudobulk profiles, and the induced changes were measured in the reconstructed expression profiles relative to 1) the control cell profiles and 2) the mean expression profile across knock-down perturbations (perturbed mean, PM) (**Figure 6**). The first test conveys the idea of absolute transcriptional changes, while the second corrects for the systematic bias in experiments in which all KDs induce similar transcriptional shifts [24]. Viñas-Torné et al. reported RPE1 cells to have a higher degree of systemic variation than K562 cells [24].

**Figure 6.**
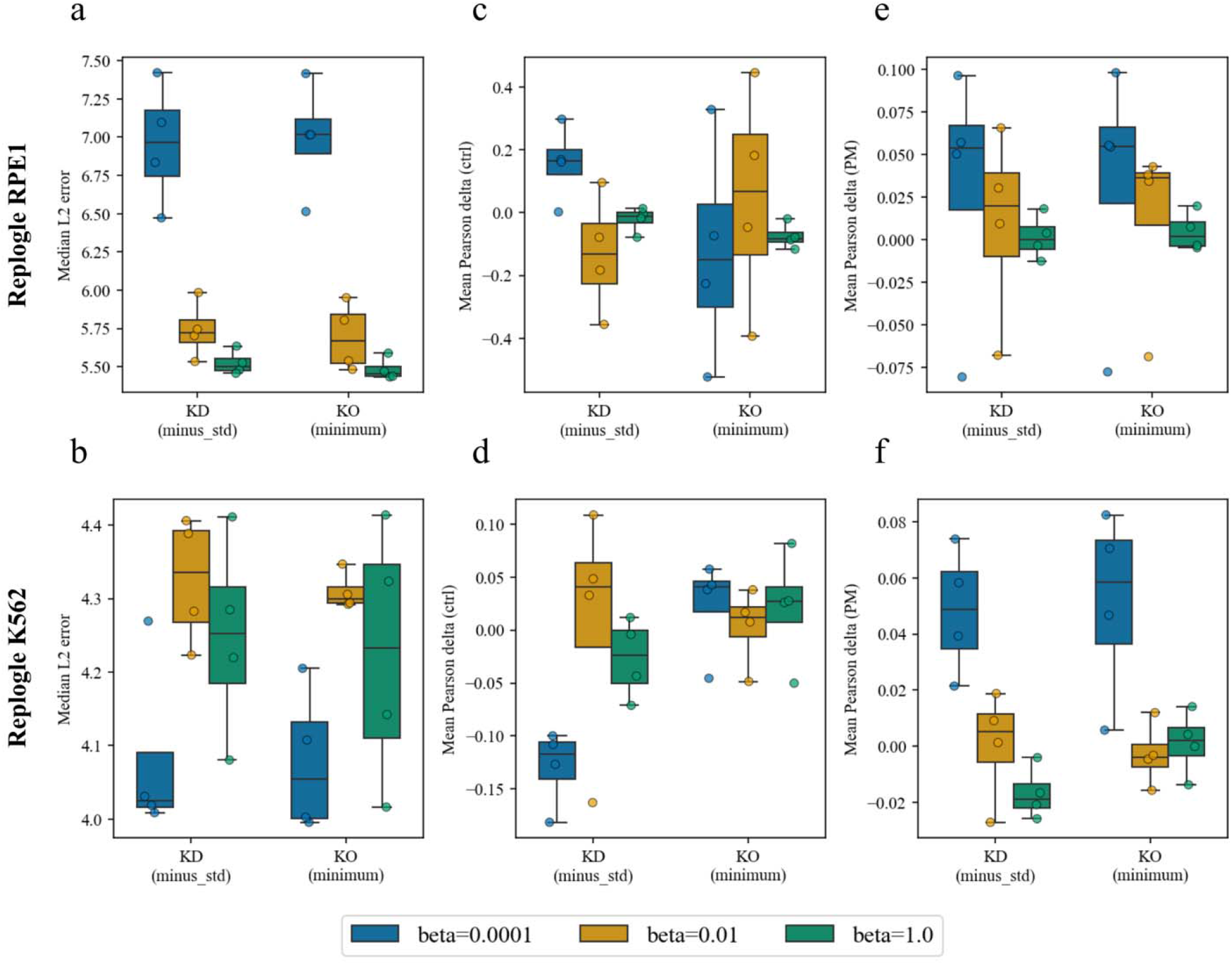
MOVE’s analyses on single cell CRISPRi data reveal shortcomings when applying *in silico* perturbations to a VAE’s input. Low-beta (=0.0001) means low regularization of the latent space, i.e. a small weight on the KLD term with a gaussian prior when training MOVE (**Methods**). **a**,**b**. Median L2 test error across 4 folds under different regularization schemes. **c, d**. Mean Pearson delta measured as the correlation between predictions and observations after subtracting the expression in the control condition. **e, f**. Mean Pearson delta measured as the correlation between predictions and observations after subtracting the mean expression profile across perturbations. Overfitting is observed at lower regularization regimes (*β*={0.0001,0.01}, blue and brown) as a high variability in prediction performance across folds. A posterior collapse is observed at high regularization (*β* = 1, green) the model predicts the perturbation mean across training samples.

Mean errors for MOVE predictions (L2) across regularization regimes ranged from 6.17 to 8.68 (RPE1) and 4.28 to 4.48 (K562) on test set genes shared across all methods (159 genes for RPE1, 149 for K562) (**Supplementary Figure 16, Supplementary Table 6, Methods**). These values were comparable to those of the other models benchmarked in a single gene KD setting by Ahlmann-Eltze et al., including foundation models such as scGPT (L2_RPE1_=6.88, L2_K562_=4.41) graph-based models like GEARS (L2_RPE1_=6.33, L2_K562_=4.04), and their proposed bilinear baseline (L2_RPE1_=5.84, L2_K562_=4.04) (**Figure 6a,b, Supplementary Figure 16**).

Pearson delta correlation means for MOVE predictions ranged from −0.178 to 0.119 (RPE1) and −0.107 to 0.022 (K562) under different regularization regimes and perturbational schemes (**Supplementary Figure 17, Supplementary Table 6, Methods**). Like CPA (Pδ_RPE1_=0.053, Pδ_K562_=0.027), the other VAE-based approach introducing perturbations in the latent space, MOVE underperformed all non-VAE-based approaches: scGPT (Pδ_RPE1_=0.642, Pδ_K562_=0.311), GEARS (Pδ_RPE1_=0.697, Pδ_K562_=0.378), the mean baseline (Pδ_RPE1_=0.739, Pδ_K562_=0.410) and the bilinear model (Pδ_RPE1_=0.744, Pδ_K562_=0.436)(**Figure 6c,d, Supplementary Figure 17**).

Expression changes after a KD were more pronounced in RPE1 cells than in K562 cells, and this higher variance in the data was captured by the trained models (**Supplementary Figures 18**,**19**). As described in previous sections, the magnitude of the predicted changes was smaller than that observed. This was particularly the case for models trained under a low regularization regime (β=1e-4) (**Supplementary Figures 18, 19**). Importantly, small predicted changes were not unique to MOVE but were also observed in the baseline proposed by Ahlmann-Eltze (**Supplementary Figure 20**). Since the Pearson delta correlation is rank-based and the L2 distance is dominated by a large number of genes whose expression barely changes after perturbation, neither metric penalize models for predicting small effect sizes.

A low regularization (*β* =0.0001) led the model to a better prediction accuracy when comparing with the perturbation mean (**Figure 6e-f**). However, this was at the expense of a higher degree of overfitting, seen as a wider spread of both loss and accuracy metrics on unseen samples across data folds (**Figure 6, blue boxes**). Higher overfitting at low regularizations was observed for different model initializations, different model sizes, and across perturbation magnitudes (**Supplementary Figures 21-23**). When increasing the regularization (={0.01, 1}), we observed a posterior collapse: cosine similarities reached the maximum while individual gene pearson correlations dropped to zero (**Supplementary Figures 24-27**). This apparent inconsistency is explained by the model predicting the mean perturbed profile while the latent space losing all sample discriminating information. As a consequence, the comparison with mean effect across perturbations (PM) became zero (**Figure 6e-f**).

In summary, we extended the use of our *in silico* perturbation paradigm in VAE-based frameworks to a new domain, namely the genetic perturbations in the context of virtual cells. Using Ahlmann-Eltze as a reference, we compared MOVE’s performance with that of alternative methods including GEARS, CPA, scGPT and their baseline linear models. These findings highlight the limitations of current VAE-based approaches when trying to infer the impact of genetic perturbations in the cell’s transcriptomes, especially when predicting the resulting cell states and not only the direction of the changes.

## Discussion

Autoencoders, in particular Variational AutoEncoders (VAEs), are powerful tools for integrating complex biomedical data [6,10,11,25] and are often used for perturbation-based downstream analyses [10,11,26]. However, their “black box” nature and potential performance benefits are often debated [11,15,17,27,28]. In this work we aimed to provide an intuitive, mechanistic perspective on the behavior of our multimodal VAE framework MOVE [10] when 1) integrating multimodal biomedical data and 2) inferring the downstream consequences of perturbing the integrated modalities. Our findings can be summarized as follows.

First, the inner functioning of the models changes depending on the nature and properties of the dataset at hand. The model’s ability to learn and represent data is shaped by factors like feature entanglement (e.g., correlations), multicollinearity from low sample-to-feature ratios, and data sparsity [14]. This dependency highlights the need for synthetic benchmarks that can accurately capture the properties of the real-world data they aim to model.

Furthermore, the model’s objective function deeply shapes the latent space organization of the samples. In a low-regularization regime (β → 0), the VAE behaves like a simple autoencoder, separating samples into distinct clusters based on categorical labels to improve reconstruction accuracy. The resulting latent space can become fragmented, with large unexplored regions between clusters. This fragmentation increases the risk of overfitting, especially with highly categorical data like genotypes. *In* silico perturbations may then push sample representations into unmapped regions, producing reconstructions that lack biological plausibility. A critical finding is that while the model can correctly infer the *direction* of change after a perturbation (e.g., increasing cholate levels in dysbiosis), the *magnitude* of this change is often small. A perturbed sample may not fully transition to its target cluster in the latent space, and its reconstructed categorical labels may even revert to the original state. The fact that perturbed samples may revert to the original categorical label urges caution when interpreting in silico perturbations as faithful biological simulations. We consider this one of the most important findings of this manuscript.

We note that this behavior is not unique to MOVE but widespread across perturbation-oriented models. In fact, standard evaluation metrics can be insensitive to the overall scale of the predicted perturbation effect. Error metrics such as the L2 distance penalise gene-level deviations, but most genes can be minimally affected by any single perturbation and hence predicting the mean perturbed profile is already a strong baseline. The Pearson delta correlation, by design, is invariant to amplitude and rewards only the correct ranking of gene-level changes. The bilinear model of Ahlmann-Eltze et al. illustrates this: its predictions concentrate on the bias term (MAD ≈ 0), yet it performs competitively on both metrics. This suggests that current benchmarks reward approximating the changes after perturbation but do not capture how meaningful the predicted perturbed state actually is.

Strategies to mitigate latent space fragmentation, such as increasing regularization (higher β) or up-weighting continuous data modalities, come at a cost, namely a drop in reconstruction accuracy. Since perturbation analyses can depend on high-quality baseline reconstructions, this trade-off directly undermines their reliability. The problem is compounded by our finding that latent space dynamics are local, not global. Our work challenges the direct application of simple vector arithmetic (as seen in language models [29–31]) to multimodal latent spaces. The effect of a perturbation is not a single, global vector; instead, it depends on the sample’s local position in the latent space. This finding aligns with recent work showing that autoencoders learn non-linear vector fields [32].

While our MOVE framework leverages VAEs to compress high-dimensional, multi-modal biological data into a latent space, alternative representation learning strategies emphasize the non-Euclidean topologies inherent to complex biological systems through the use and interpretation of graphs [17,33]. Recent studies utilizing Heterogeneous Information Networks (HINs) and Graph Neural Networks (GNNs) explicitly preserve the structural priors of known interactions by mapping connections through robust frameworks like STRING network embeddings and the multiscale interactome [34,35]. Rather than simulating physiological variations via *in silico* generative perturbations to infer entanglement, these models treat association discovery as a discriminative link prediction task. This topological approach has proven highly effective for predicting complex miRNA-disease associations [36,37] and advancing zero-shot drug repurposing for diseases that lack existing treatments [38,39]. Despite having divergent architectures, these methodologies converge on the critical importance of localized context. For instance, GNNs rely heavily on extracting neighborhood-level structural motifs to capture non-linear interactions and prevent biologically implausible predictions. Ultimately, integrating explicit network topologies with the capabilities of VAEs might represent a promising avenue for future multi-omics integration.

In conclusion, integrating diverse biomedical data is a challenging task that requires a deep understanding of both the data and the models internal behavior. A better mechanistic interpretability of the models helps identify pitfalls, builds trust between experimental and modeling communities, and helps generating new research questions. Future research should continue to explore how perturbations propagate through more complex architectures to better separate true biological signals from model artifacts [40]. We hope this work provides a valuable perspective for researchers using VAEs for biomedical data integration and *in silico* perturbational analyses.

## Methods

### Data obtention and preprocessing

#### Synthetic datasets

Synthetic datasets emulated multimodal readouts from samples across ***N* = 5**,***000*** samples across ***P* = 152** features organized into two continuous blocks (Continuous_A: 50 features; Continuous_B:100 features) and two categorical features. Each sample *x* ∈ ℝ ^***P***^ was drawn from a multivariate Gaussian *x* ∼ **𝒩** (***μ, Σ***) where ***μ*** was a structured mean vector and *Σ* was a sparse covariance matrix with sparsity ***α***_*cov*_ = **0.99**

#### Feature configuration

Synthetic datasets consisted of *P* =152 features organized into four blocks: Continuous_A (*P*_*A*_ = features), Continuous_B (*P*_*B*_ = features) Categorical_A (*P* _*catA*_ =1feature), and Categorical_B (*P* _*catB*_ =1feature). Features were indexed as *p* = 1,…, *P* in this block order.

#### Mean vector

The mean of each feature was determined by a structured sinusoidal profile. For a feature at position within its block, the mean was:

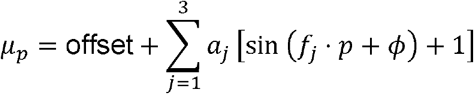

where offset, *a*_*j*_ (coefficients), *f*_*j*_ (frequencies), and *ϕ* (phase) were block-specific parameters. This produced feature means in the range ∼1350–1488 for Continuous_A, ∼467–520 for Continuous_B, and ∼10–20 for the categorical blocks.

#### Covariance matrix

The covariance matrix **Σ** ∈ ℝ ^*P* × *P*^was generated using `sklearn.datasets.make_sparse_spd_matrix` with sparsity parameter *α*_*cov* =_0.99, coefficient range [0,1], and unnormalized diagonal (`norm_diag=False`). This function constructs a sparse symmetric positive-definite matrix by first generating a sparse matrix with the specified fraction (1 − *α*_*cov*_ = 0.01) of non-zero off-diagonal entries, then ensuring positive-definiteness through inversion of a sparse precision matrix. The random seed was fixed at 1 for reproducibility.

To create a high-correlation regime (condition III), the feature set was split into two halves ℱ _1_ = {1,…,76} and ℱ _2_ = {77,…,152}. Each feature was redefined as: *k* ∈ ℱ _2_ was redefined as:

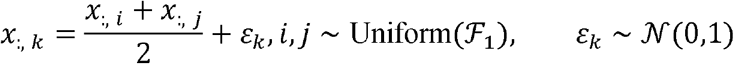

where *ε* _*k*_ was a single scalar offset per feature. Categorical features were subsequently binarized by thresholding at the feature mean. Ground-truth associations were defined as feature pairs with absolute Pearson correlation exceeding *r* _thres_ = 0.02.

Because ℱ _2_ included the categorical feature indices (*p* = 151,152), these categorical features also inherited linear dependencies on first-half features before binarization (see below).

#### Binarization of categorical features

After the correlation injection step, the *P*_cat_ = 2 categorical features (the last columns of the dataset) were binarized: for each categorical feature, values above the feature mean were set to 1, and values at or below the mean were set to 0.

#### Preprocessing

No scaling or log-transformation was applied during dataset generation. These steps (log_2_(1 + *x*) transformation followed by z-score normalization to zero mean and unit variance) were performed downstream in the MOVE data preprocessing pipeline.

#### Preprocessing of IBDMDB data

Data was obtained from the IBDMDB database (https://ibdmdb.org/), and all data processing steps are detailed in the main jupyter notebook of the project, which can be found <u>here</u>. Metabolomics, metatranscriptomics and metagenomics data were log_2_(1+x) transformed to reduce their dynamic range and posteriorly z-scored normalized. Metagenomics and metatranscriptomics features with more than 50% of zero counts across samples were discarded from the analyses. Only metabolites with common identifiers were included to ease interpretability. In MOVE, NA values would be imputed by 0 after feature scaling, which would correspond to an imputation by the mean. In order to prevent MOVE from imputing measurements with zero counts by the corresponding feature mean across samples, which would not convey the idea that that metabolite / species was not present in the sample, zeros were not transformed to NAs.

#### Preprocessing of Perturb-seq data from Replogle et al

Data from the perturb-seq analyses on K562 and RPE1 cells was obtained following the procedure described in Ahlmann-Eltze et al., i.e. we obtained the essential dataset from GEARS [13,17,18]. The data was already log-transformed, so we did not log_2_(1+x) transform it. To facilitate the comparison with the other models, we did not z-score transform the data either, i.e. we computed pseudobulks for each gene knock-down (KD) and provided them directly as inputs to MOVE. This allowed us to compute perturbation effects directly in expression space and not have to invert the z-score mapping but influenced the scale of the standard deviation and the minimum feature values for the cohort.

The GEARS preprocessing pipeline applies a highly variable gene (HVG) selection (∼5,000 genes). Because MOVE perturbs genes by modifying their expression values in the input, only perturbation targets that are also present in the HVG set can be evaluated: 412 out of 1,092 target genes in K562 (37.7%) and 651 out of 1,543 in RPE1 (42.2%). The remaining target genes fall outside the HVG set and cannot be perturbed *in silico*. When restricting the comparison to genes shared across all 22 methods in the Ahlmann-Eltze et al. benchmark, this overlap is further reduced to 149 genes for K562 and 159 for RPE1. We used a 4-fold cross-validation scheme on perturbation genes, partitioning the set of unique target genes into 4 non-overlapping folds (∼75% train, ∼25% test), matching the 75/25 train/test gene split used by Ahlmann-Eltze et al.

### Analysis and benchmarking

#### The MOVE framework

##### Variational AutoEncoders

Variational AutoEncoders (VAEs) are a type of artificial neural networks that use variational inference to approximate the distribution of our data p(**x**). This is achieved by maximizing the evidence lower bound (ELBO), a lower bound on the log-likelihood:

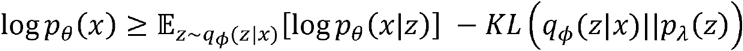

The ELBO consists of two terms, the reconstruction term (first) and a regularizer term (second). Here, we consider that our data **x** has some hidden factors **z**. The hidden factors can be obtained by sampling from a prior distribution **z ∼** p_λ_(**z**), and can be decoded to yield data samples **x ∼** p_θ_(**x** | **z**). However, we use an amortized variational posterior to generate the latent representations of our samples **z ∼** p_φ_(**z** | **x**). These distributions are parametrized using neural networks: a stochastic encoder with parameters {φ}, which yields means (μ) and standard deviations (σ) for each sample’s latent representation; a stochastic decoder with parameters {θ} that maps latent representations back to input space, and a prior distribution with parameters {λ}.

MOVE was designed to be a flexible framework to build multimodal VAEs [4], creating fully connected networks that could have a varying number of hidden layers with an arbitrary number of nodes. The encoder was provided the concatenated inputs of all datasets for a given batch of samples. Batch normalization and dropout were performed after each fully connected hidden layer, for which we used LeakyReLUs as the activation function [41]. The final models for this paper and the original MOVE used one hidden layer only [10]. The encoder network ended in a linear latent layer with N_L_ nodes encoding the means (*μ*) and N_L_ nodes for the logarithms of the variances (log *σ*^2^) with their respective biases. During training, the reparameterization trick (*z* = *μ σ · ϵ*) was performed before propagating the representations of the samples through the decoder, where *ϵ* was drawn from a normal Gaussian. The decoder then ended with a linear layer, from which we mapped back the representations to input space.

##### Model training

The network was trained to optimize a Mean Squared Error (MSE) loss for the continuous variables and a Categorical Cross Entropy (CCE) for categorical variables. The latter was implemented by obtaining log-probabilities of the different categories by applying a log-softmax to the output nodes corresponding to categorical variables, which were then subject to a Negative Log Likelihood loss, as implemented in Pytorch [42]. A Normal Gaussian was used as the prior distribution of latent variables, which we enforced by adding the Kullback-Leibler divergence (KLD) term to the loss.

If we assume that continuous features follow a Gaussian distribution with mean μ and fixed variance σ^2^, we have

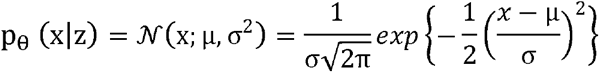

Minimizing the negative log-likelihood is, in this case, equivalent to minimizing the mean squared error of the predictions when sigma is constant:

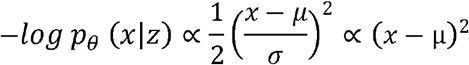

In a similar fashion, categorical variables follow a categorical distribution *p*_*θ*_ (*x*|*z*) = *Cat* (*x*;*p*_1_, …,*p*_*K*_) where each class or category has a probability assigned *p*_*i*_. The negative log-likelihood in this case, corresponding to the categorical cross entropy loss, is:

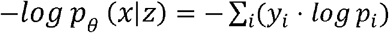

The contribution of missing features to the loss was removed and did not contribute to backpropagation. For categorical variables, this was achieved by assigning a dummy index (-1) to zero vector features, which was then ignored when computing the loss. For continuous variables, outputs where the corresponding input values were filled with zeroes were also set to zero. This yielded a zero contribution to the mean squared error computation for that node. Since the gradient of the MSE loss for a given node is proportional to the value difference between the predicted output and the target, i.e. the input, the gradients of the loss towards the node were also zero.

##### Loss function

Let 𝒞 denote the set of categorical datasets and 𝒟 the set of continuous datasets. The total loss for a mini-batch of *B* samples was:

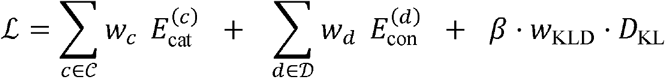

Each term is defined below.

##### Categorical reconstruction error

For a categorical dataset *c* containing *P*_*c*_ categorical features, each with *K*_*c*_ classes, the error was computed as:

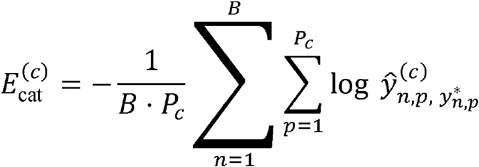

where 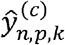 are the softmax output probabilities for sample *n*, feature *p*, and class *k*, and 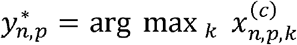 is the ground-truth class index derived from the one-hot input x. Features with all-zero one-hot vectors (missing data) were assigned a dummy index of −1 and excluded from the loss computation.

##### Continuous reconstruction error

For a continuous dataset *d* containing *p*_*d*_ features:

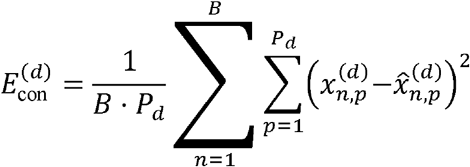

Prior to this computation, reconstructed values 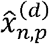 were set to zero wherever the corresponding input 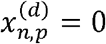, effectively masking zero-valued entries so that they contributed neither to the loss nor to the gradient. This served as a missing-data handling strategy.

##### KL divergence

A standard Gaussian prior *p*(*z*) = 𝒩 (**0**,**I**) was imposed on the *N*_*L*_-dimensional latent space. The KL divergence was:

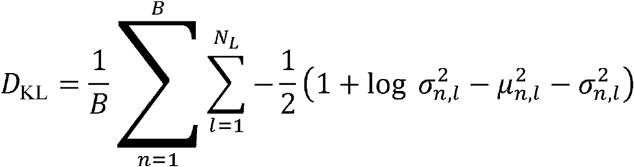

where the encoder network outputs *μ* _*n,l*_ and 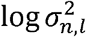 directly (i.e., the log-variance, not the log-standard-deviation).

##### Dataset weights

Each *w*_*c*_ and *w*_*d*_ is a non-negative scalar hyperparameter controlling the relative contribution of its dataset to the total loss. These weights are unconstrained (they need not sum to one), default to 1, and should be set in a dataset-dependent fashion. For the IBDMDB analyses, the continuous dataset weights are denoted *w*_mtx_(metatranscriptomics), *w*_mgx_ (metagenomics), and *w*_mbx_ (metabolomics).

When no per-dataset weights are specified, the categorical term reduces to the unweighted mean across datasets: 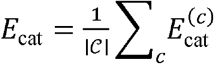, and the continuous term is computed as a single pooled MSE over all continuous features: 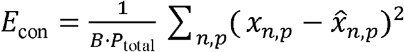.

##### KLD weight schedule

The effective weight on the KL term is the product of two factors: *β*, a fixed hyperparameter, and *w*_KLD_ = *i* _*step*_ ·*β* /*N* _steps_, a warm-up schedule that linearly increases from 0 to *β*over a predefined number of training steps. In the IBDMDB association analyses, *β* = 10^−4^, meaning the regularization term was effectively negligible.

Models were trained using the Adam optimizer [43]. Batch dilation was introduced to increase progressively the batch size. Finally, latent representations at inference time were given by the *μ* vector.

##### Hyperparameter tunning

We did not perform hyperparameter tuning for the latent space analyses, as our goal was to explore the VAE’s behavior under different hyperparameter regimes. For identifying associations in the IBDMDB dataset, however, we performed hyperparameter tuning to maximize reconstruction quality, measured as the cosine similarity between input and output vectors for continuous data. We explored combinations of batch size = {16,32}, N_hid_ = {256, 512} and N_lat_ = {8,32,64}, while keeping the number of epochs fixed at 400, model refits at 24, learning rate at 1e-4, and β at 1e-4. These fixed values ensured proper baseline reconstructions. The smaller batch size consistently produced better reconstructions, and quality increased with N_lat_ up to 32 before stabilizing. As hidden layer dimensionality had little impact, we proceeded with (batch size, N_hid_, N_lat_) = (16,256,32). Notably, both categorical variables (IBD status and Dysbiosis) were perfectly reconstructed regardless of the hyperparameters (**Supplementary Figure 6**).

Empirically, setting all dataset weights (*w* _cat,_ *w*_con_) to one while normalizing for feature number and batch size boosted the contribution of the categorical variables to the loss. Continuous dataset weights were set to 10 to show the impact of continuous dataset boosting in the synthetic scenario, and they were set to 2 when identifying associations in the IBDMDB dataset when boosting the contributions of continuous data modalities. 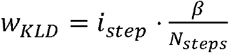 was used as the weight assigned to the KLD term, controlled by a parameter *β* defining the maximum attainable *w*_*KLD*_ and starting from 0. A warm-up procedure for the KLD increased its weight to a higher fraction in a number of steps, where each step was a predefined epoch.

When identifying associations, *β* was set to the original 1e-4, i.e. the regularization term was in practice omitted to get reliable baseline reconstructions. Under this low regularization regime, the reconstruction accuracies of hold-out validation samples were stable across multiple model hyperparameter combinations, so we kept the simplest well-performing model. Specific details of the hyperparameters of the different models used in the paper can be found on the configuration files in the Github repository of the project.

##### Feature importance, SHAP-based analysis

The adaptation of the SHAP algorithm presented in [10] was used to define the importance that the model attributed to each input feature. The input dataset **X**, a matrix with N_samples_ rows and N_features_ columns, served as a reference and was encoded into its latent representation **z**, with N_samples_ rows and N_latent_ columns. A perturbed dataset **X’**, where the feature of interest had been substituted in all samples for the missing value (i.e. 0), was encoded into its latent representation **z’**. The induced movement of the samples in latent space was then computed as the Euclidean distance **d** between both encodings, i.e. **d** = **z’ - z**. Finally, the overall movement of each sample was obtained by adding the movements in all latent components, as a sum of all elements in each row. This protocol was repeated for each feature, one by one, across all features. An ensemble of 24 models with different hyperparameters was used when performing SHAP analysis of the IBDMDB data, aiming to identify features that were important for the models regardless of specific hyperparameter choices. These were all possible combinations of models with β = {0.01, 0.0001}, N_hid_ = {64,256,512}, N_lat_= {16,32} and weights on the loss of all continuous data modalities w_mtx_, w_mgx_,w_mbx_ = {1,2}.

##### Perturbations of continuous inputs

The proposed pipeline was conceptually an extension to the SHAP protocol. A first perturbational approach consisted on substituting, in all samples, the value of the feature to perturb for the minimum or maximum found in the cohort. Note that this was done after standardization to zero mean and unit variance for each feature, and therefore depending on the original distribution of the feature and the number of samples this could yield arbitrarily large or small values. The second approach consisted on adding or subtracting one standard deviation to the feature to perturb in all samples. This allowed us to keep the previous context of the feature in a given sample. The small quantity to add or subtract was chosen to one standard deviation of the value distribution, which describes its scale of variation.

##### Finding associations between variables

The MOVE t-test and Bayes inspired algorithms were developed in Allesøe et al. [10], and are described in detail in the following sections. Here we proposed the Kolmogorov-Smirnov (KS) distance, 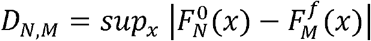, to compare empirical distributions of baseline 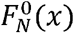 and perturbed 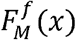 feature reconstructions, where N=M. We used the median of signed distances (*D*_*N*_) across refits to minimize outlier impact. An association was identified if the null hypothesis, namely that baseline and perturbed reconstructions originated from the same distribution, was rejected. The rejection condition, 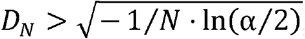, is a function of the significance threshold α. Two-sample KS tests with a small sample sizes entail a high *D*_*N*_, meaning only large distributional shifts are considered to be significant associations (**Supplementary Figure 14**). The number of samples was therefore a significant limiting factor. The sign of the KS score indicates the directionality of the association; for instance, a positive perturbation on a positively related variable results in a negative KS score (**Supplementary Figure 4e**).

##### Finding associations with MOVE t-test

The t-test approach was presented in the original MOVE paper. In this method, the expected average change in the reconstructions when passing the original and perturbed data through the model a number of times was used to determine the significance of the associations between variables. In this work, we simply extend the perturbations in the input to the continuous domain, but the measurements in the output remain the same. A Student’s t-test, as implemented in Scipy [44], was performed between the reconstructions of unperturbed samples and the reconstructions of the samples for which a given feature was perturbed. The obtained p-values were then Bonferroni-corrected for each perturbation independently, and we set the significant threshold as adjusted p < 0.05. The analysis was repeated ten times, i.e. the models had 10 refits, for four different latent sizes. The associations were accepted only if they were significant in three out of the four possible latent sizes and at least half of the refits. Thus, the final p-values were the averages over the ten refits for four latent sizes, yielding 40 two-sided Bonferroni-corrected t-tests. The reported effect sizes for this method corresponded to the magnitude of the obtained p-value, i.e. -log_10_(p-value).

##### Finding associations with Bayes decision theory

This approach was also described in MOVE [10], and here was extended for the perturbations of continuous variables. The method was inspired by Bayesian decision theory and the approach follows the ideas presented in [45], an application for single cell variational inference, and [46]. Given a fixed VAE architecture, we trained the model a number of times *N*_*refits*_, which we called refits. For VAE refit *i*, sample or measurement *n* and target feature *t* we obtained the reconstructions of the sample at the model’s output in two conditions: when using the baseline samples as inputs 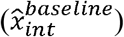 and also for samples where a feature of interest *f* was perturbed 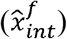. We then measured the differences between reconstructions 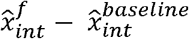, averaged them across refits, and computed the fraction of samples *q*_*tf*_ in which the averaged difference was positive. *q*_*tf*_ was used to compute a score for each perturbed-target feature pair:

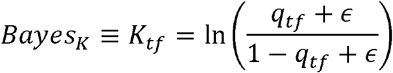

Where ϵ = 1 · 10^−8^ was added for numerical stability. The score was finally introduced into a logistic function 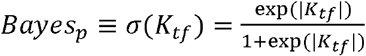 to obtain a probability-like value, bound between 0 and 1, for each possible pair of perturbed and target features. The values of *Bayes* _*K*_ and *Bayes* _*p*_ for any arbitrary fraction of samples *q*_*tf*_ are shown in **Supplementary Figure 15**. *Bayes* _*p*_ was finally used to rank the pairs of perturbed and target features, and associations were considered significant until the cumulative evidence across accepted pairs reached a significance threshold 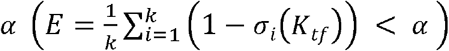, where *i* is the index of the association in the ranked list. In practice, a high number of associations were assigned the highest probability (*Bayes*_*p*_ =1), and therefore the contribution of the first associations to the cumulative evidence was driven by *ϵ*.

##### Benchmarking MOVE with other perturbational approaches on Replogle et al. data

We trained MOVE in a 4-fold validation setting to compress and reconstruct the pseudobulk profiles on RPE1 and K562 Perturb-seq data from Replogle et al. [13]. Once the model had been trained across knock-downs, *in silico* perturbations were applied in the input by lowering the expression values for the perturbed gene. Perturbations were applied to the control pseudobulk profile on unseen test perturbations. Minimum feature values and standard deviations per feature were computed on training samples. Inference was run on a number of MOVE models under different regularization regimes (*β* ∈ {0.0001,0.01,1}) and perturbation magnitudes (KD i.e. minus std, KO i.e. minimum value substitution) scenarios. All metrics are computed on held-out (test-split) perturbation conditions and restricted to the top 1000 most expressed genes, following Ahlmann-Eltze et al. We compared MOVE prediction performances against the performance metrics L2 and r2_delta for single gene KDs reported in panel C in the <u>Source Data file for Figure 2 in Ahlmann-Eltze et al</u>.

##### Bilinear perturbation model by Ahlmann-Eltze et al

We reproduced the baseline bilinear perturbation model on pseudobulk RNA-seq profiles presented by Ahlmann-Eltze et al. in [18]:

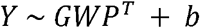

With *Y* being the expression matrix (*n*_*genes*_ × *n*_*perturbations*_), *G* being the gene embedding obtained as a PCA on *Y*_*trian*_(*n*_*genes*_ × K, K =10),*P* representing the perturbation embedding chosen to be the rows of *G* matching the perturbed genes, and *b* being the bias or mean baseline obtained as the mean across perturbations, i.e. row means of *Y*_*train*_. The weight matrix *W* can be solved analytically via Ridge-regularized normal equations:

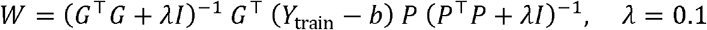

The goal was to quantify the magnitude of the predicted changes, i.e. the deviation from the mean provided by the term on the equation, shown in **Supplementary Figure 20**.

##### Evaluation metrics

###### Continuous Reconstruction Accuracy (Cosine Similarity)

Measures the angular similarity between the original continuous feature vector (*x*) and its reconstruction 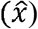 to quantify reconstruction quality. Obtained for each continuous data modality separately.

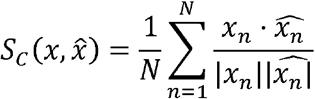

###### SHAP (Latent Space Impact)

Quantifies the importance of a feature by measuring the average Euclidean distance a sample’s representation moves in the latent space (*z*) upon feature perturbation.

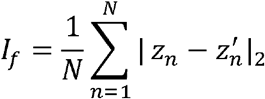

###### Bayesian Association Score (Bayes K)

A log-odds ratio used to identify significant associations, calculated based on the fraction of samples (*q*_*tf*_) with positive reconstruction changes for a target feature *t* given perturbation of feature *f*.

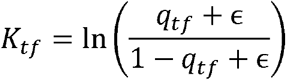

###### Bayesian Association Probability (Bayes p)

A normalized value between 0 and 1 derived from the Bayes K score using a logistic-like transformation to provide a probability-like significance measure.

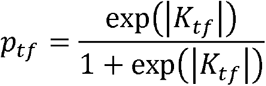

###### Cumulative Evidence (E)

A ranking metric that selects significant associations by summing the uncertainty (1 - p) across accepted pairs until a specified threshold *α* is reached.

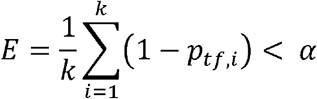

###### Kolmogorov-Smirnov (KS) Distance

A non-parametric metric measuring the maximum distance between the cumulative distribution functions of baseline and perturbed reconstructions to capture distribution shifts.

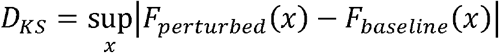

###### L2 Prediction Error

Measures the overall prediction accuracy as the Euclidean distance between the predicted expression profile 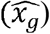 and the observed ground truth profile ***x***_*g*_ following a perturbation of gene *g*. The evaluation is performed on the 1000 most highly expressed genes in the control condition following Ahlmann-Eltze et al. [18] :

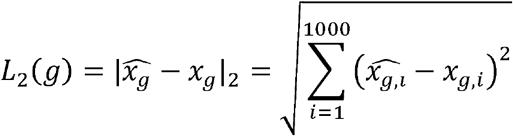

###### Pearson Delta vs Control ρ_ctrl_)

The correlation between predicted and observed expression changes relative to the unperturbed control, assessing the model’s ability to capture the pattern and direction of differential expression.

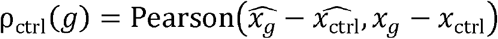

###### Pearson Delta vs Perturbed Mean (ρ_PM_)

A metric measuring the correlation of predicted and observed deviations relative to the mean knock-down profile, testing for gene-specific prediction accuracy beyond average effects.

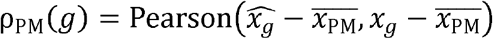

## Declarations

### Ethics approval and consent to participate

Not applicable.

### Consent for publication

Not applicable.

### Clinical trial number

Not applicable.

## Availability of data and materials

The synthetic datasets presented in this manuscript can be created using the code provided at the Github repository of the project. IBDMDB raw data was obtained from the IBDMDB database (https://ibdmdb.org/). Data from Replogle’s perturb-seq analyses on K562 and RPE1 cells was obtained following the procedure described in Ahlmann-Eltze et al., i.e. we obtained the essential dataset from GEARS [13,17,18]. All data processing steps are detailed in the main jupyter notebook of the project, which can be found <u>here</u>. Code and configuration files to create and run the MOVE models used in this manuscript can be found at the Github repository of the project: https://github.com/RasmussenLab/VAEs_for_biomedical_data_integration.

A tutorial notebook covering the main findings presented in this manuscript is also available: https://colab.research.google.com/github/RasmussenLab/VAEs_for_biomedical_data_integration/blob/main/scripts/Tutorial_VAEs_for_biomedical_data_integration.ipynb

MOVE can be downloaded as a pip package as move-dl.

## Authors’ contributions

S.R. conceived the study and guided the analysis. M.P.A. performed the analyses, wrote the code, interpreted the results, extended the framework to handle continuous perturbations and proposed the KS method. R.H.M. developed the bayes inspired approach, wrote the code and helped in the analyses. H.W. wrote the code and helped in the analyses. All authors reviewed and approved the final manuscript.

## Acknowledgements

We would like to thank the members of the Rasmussen Lab, in particular Pau Piera, Jonas Meisner, and Leonardo Cobuccio for their feedback and fruitful discussions. Biorender images were used in **Figures 1** and **5**. We would like to thank the Novo Nordisk Foundation Center for Basic Metabolic Research and the Novo Nordisk Foundation Center for Genomic Mechanisms of Disease for their support.

## Supplementary information

**Additional file 1**. Supplementary Figures 1 to 27.

**Additional file 2**. Supplementary Tables 1 to 6.

## References

1. Altman N, Krzywinski M. The curse(s) of dimensionality. Nat. Methods 2018; 15:399–400

2. Xu C, Jackson SA. Machine learning and complex biological data. Genome Biol. 2019; 20:76

3. Moon KR, van Dijk D, Wang Z, et al. Visualizing structure and transitions in high-dimensional biological data. Nat. Biotechnol. 2019; 37:1482–1492

4. Kingma DP, Welling M. Auto-Encoding Variational Bayes. arXiv [stat.ML] 2013;

5. Allesøe RL, Nudel R, Thompson WK, et al. Deep learning–based integration of genetics with registry data for stratification of schizophrenia and depression. Science Advances 2022; 8:eabi7293

6. Tu X, Cao Z-J, Chenrui X, et al. Cross-Linked Unified Embedding for cross-modality representation learning. Advances in Neural Information Processing Systems 2022; 35:15942–15955

7. Grønbech CH, Vording MF, Timshel PN, et al. scVAE: variational auto-encoders for single-cell gene expression data. Bioinformatics 2020; 36:4415–4422

8. Nissen JN, Johansen J, Allesøe RL, et al. Improved metagenome binning and assembly using deep variational autoencoders. Nat. Biotechnol. 2021; 39:555–560

9. Líndez PP, Johansen J, Kutuzova S, et al. Adversarial and variational autoencoders improve metagenomic binning. Commun Biol 2023; 6:1073

10. Allesøe RL, Lundgaard AT, Hernández Medina R, et al. Discovery of drug–omics associations in type 2 diabetes with generative deep-learning models. Nat. Biotechnol. 2023; 41:399–408

11. Lotfollahi M, Klimovskaia Susmelj A, De Donno C, et al. Predicting cellular responses to complex perturbations in high-throughput screens. Mol. Syst. Biol. 2023; 19:e11517

12. Lloyd-Price J, Arze C, Ananthakrishnan AN, et al. Multi-omics of the gut microbial ecosystem in inflammatory bowel diseases. Nature 2019; 569:655–662

13. Replogle JM, Saunders RA, Pogson AN, et al. Mapping information-rich genotype-phenotype landscapes with genome-scale Perturb-seq. Cell 2022; 185:2559–2575.e28

14. Elhage N, Hume T, Olsson C, et al. Toy Models of Superposition. arXiv [cs.LG] 2022;

15. Bunne C, Roohani Y, Rosen Y, et al. How to build the virtual cell with artificial intelligence: Priorities and opportunities. arXiv [q-bio.QM] 2024;

16. Roohani YH, Hua TJ, Tung P-Y, et al. Virtual Cell Challenge: Toward a Turing test for the virtual cell. Cell 2025; 188:3370–3374

17. Roohani Y, Huang K, Leskovec J. Predicting transcriptional outcomes of novel multigene perturbations with GEARS. Nat. Biotechnol. 2024; 42:927–935

18. Ahlmann-Eltze C, Huber W, Anders S. Deep-learning-based gene perturbation effect prediction does not yet outperform simple linear baselines. Nat. Methods 2025; 22:1657–1661

19. Yang F, Wang W, Wang F, et al. scBERT as a large-scale pretrained deep language model for cell type annotation of single-cell RNA-seq data. Nat. Mach. Intell. 2022; 4:852–866

20. Theodoris CV, Xiao L, Chopra A, et al. Transfer learning enables predictions in network biology. Nature 2023; 618:616–624

21. Rosen Y, Roohani Y, Agrawal A, et al. Universal cell embeddings: A foundation model for cell biology. bioRxiv 2023;

22. Cui H, Wang C, Maan H, et al. scGPT: toward building a foundation model for single-cell multi-omics using generative AI. Nat. Methods 2024; 21:1470–1480

23. Hao M, Gong J, Zeng X, et al. Large-scale foundation model on single-cell transcriptomics. Nat. Methods 2024; 21:1481–1491

24. Viñas Torné R, Wiatrak M, Piran Z, et al. Systema: a framework for evaluating genetic perturbation response prediction beyond systematic variation. Nat. Biotechnol. 2025;

25. Cao Z-J, Gao G. Multi-omics single-cell data integration and regulatory inference with graph-linked embedding. Nat. Biotechnol. 2022; 40:1458–1466

26. Møller AF, Skat Madsen JG. Disentangling covariate effects on single cell-resolved epigenomes with DeepDive. bioRxiv 2025;

27. Qi X, Zhao L, Tian C, et al. Predicting transcriptional responses to novel chemical perturbations using deep generative model for drug discovery. Nat. Commun. 2024; 15:9256

28. Ahlmann-Eltze C, Huber W, Anders S. Deep learning-based predictions of gene perturbation effects do not yet outperform simple linear methods. bioRxiv 2024;

29. Mikolov T, Yih W-T, Zweig G. Linguistic Regularities in Continuous Space Word Representations. Proceedings of the 2013 Conference of the North American Chapter of the Association for Computational Linguistics: Human Language Technologies 2013; 746–751

30. Park K, Choe YJ, Veitch V. The linear representation hypothesis and the geometry of large language models. arXiv [cs.CL] 2023;

31. Pennington J, Socher R, Manning C. GloVe: Global Vectors for Word Representation. Proceedings of the 2014 Conference on Empirical Methods in Natural Language Processing (EMNLP) 2014; 1532–1543

32. Fumero M, Moschella L, Rodolà E, et al. Navigating the latent space dynamics of neural models. arXiv [cs.LG] 2025;

33. Cetin S, Sefer E. A graphlet-based explanation generator for graph neural networks over biological datasets. Curr. Bioinform. 2025; 20:840–851

34. Hu D, Schaap-Johansen A-L, Villarroel J, et al. Molecular maps of diseases from omics data and network embeddings. bioRxiv 2025;

35. Ruiz C, Zitnik M, Leskovec J. Identification of disease treatment mechanisms through the multiscale interactome. Nat. Commun. 2021; 12:1796

36. Zhao B-W, Su X-R, Yang Y, et al. A heterogeneous information network learning model with neighborhood-level structural representation for predicting lncRNA-miRNA interactions. Comput. Struct. Biotechnol. J. 2024; 23:2924–2933

37. Zhao B-W, He Y-Z, Su X-R, et al. Motif-aware miRNA-disease association prediction via hierarchical attention network. IEEE J. Biomed. Health Inform. 2024; 28:4281–4294

38. Zhao B-W, Su X-R, Hu P-W, et al. A geometric deep learning framework for drug repositioning over heterogeneous information networks. Brief. Bioinform. 2022; 23:

39. Huang K, Chandak P, Wang Q, et al. A foundation model for clinician-centered drug repurposing. Nat. Med. 2024; 30:3601–3613

40. Balestriero R, Lecun Y. How Learning by Reconstruction Produces Uninformative Features For Perception. Proceedings of the 41st International Conference on Machine Learning 21--27 Jul 2024; 235:2566–2585

41. Andrew L. Maas, Awni Y. Hannun, Andrew Y. Ng. Rectifier Nonlinearities Improve Neural Network Acoustic Models.

42. Paszke A, Gross S, Massa F, et al. PyTorch: An Imperative Style, High-Performance Deep Learning Library. Advances in Neural Information Processing Systems 2019; 32:

43. Kingma DP, Ba J. Adam: A Method for Stochastic Optimization. arXiv [cs.LG] 2014;

44. Virtanen P, Gommers R, Oliphant TE, et al. SciPy 1.0: fundamental algorithms for scientific computing in Python. Nat. Methods 2020; 17:261–272

45. Lopez R, Regier J, Cole MB, et al. Deep generative modeling for single-cell transcriptomics. Nat. Methods 2018; 15:1053–1058

46. Lopez R, Boyeau P, Yosef N, et al. Decision-Making with Auto-Encoding Variational Bayes. arXiv [stat.ML] 2020;

