## Supplementary Figures for "On the use of variational autoencoders for biomedical data integration"

**Author List Footnotes**

Marc Pielies Avellí

Ricardo Hernández Medina

Henry Webel

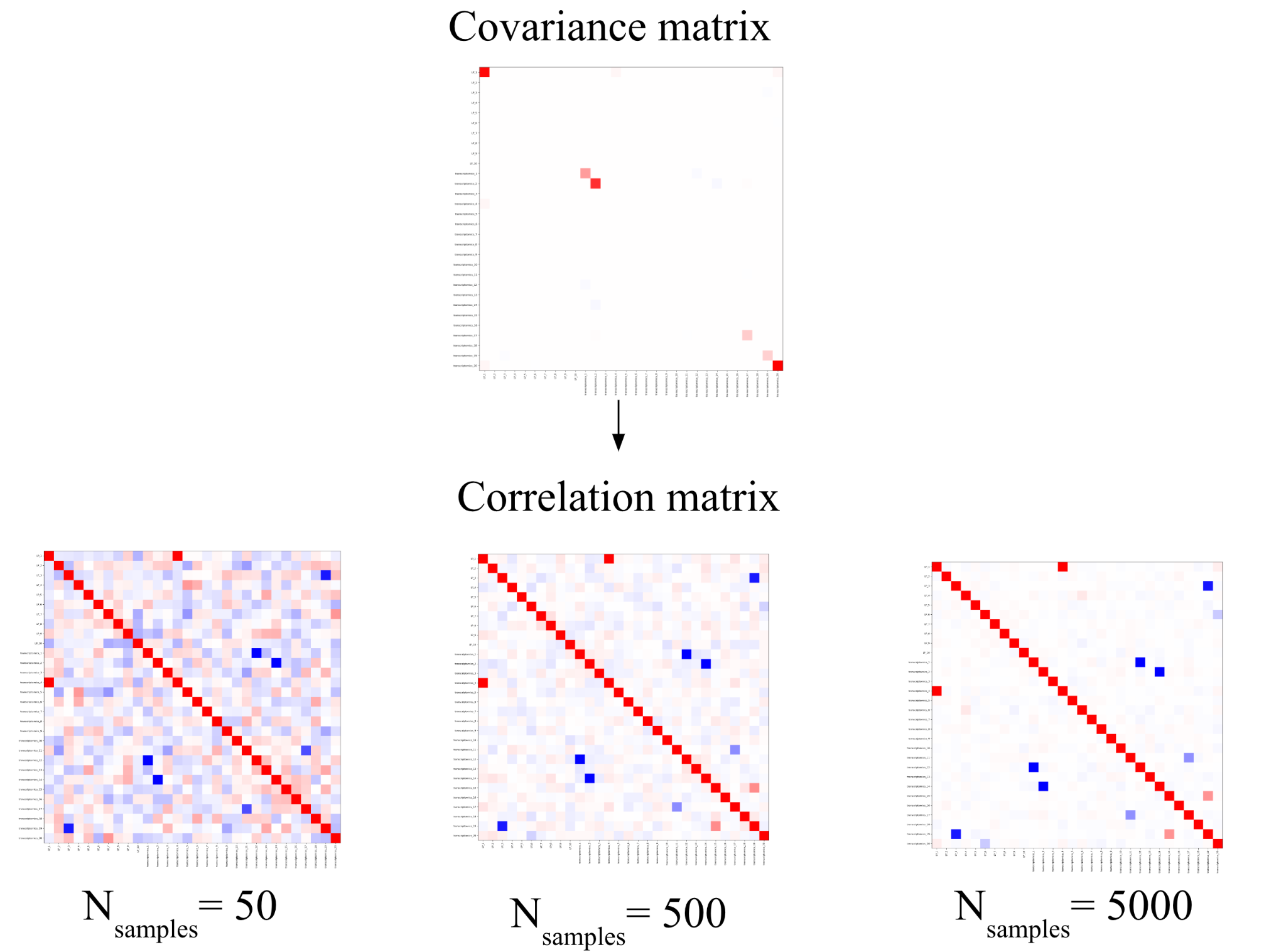

**Supplementary Figure 1. Ground truth relations between variables in a synthetic dataset.** Given a covariance matrix and a mean vector, we can obtain multimodal profiles by drawing samples from the corresponding multivariate gaussian. Background correlations emerge as artifacts under low sampling regimes.

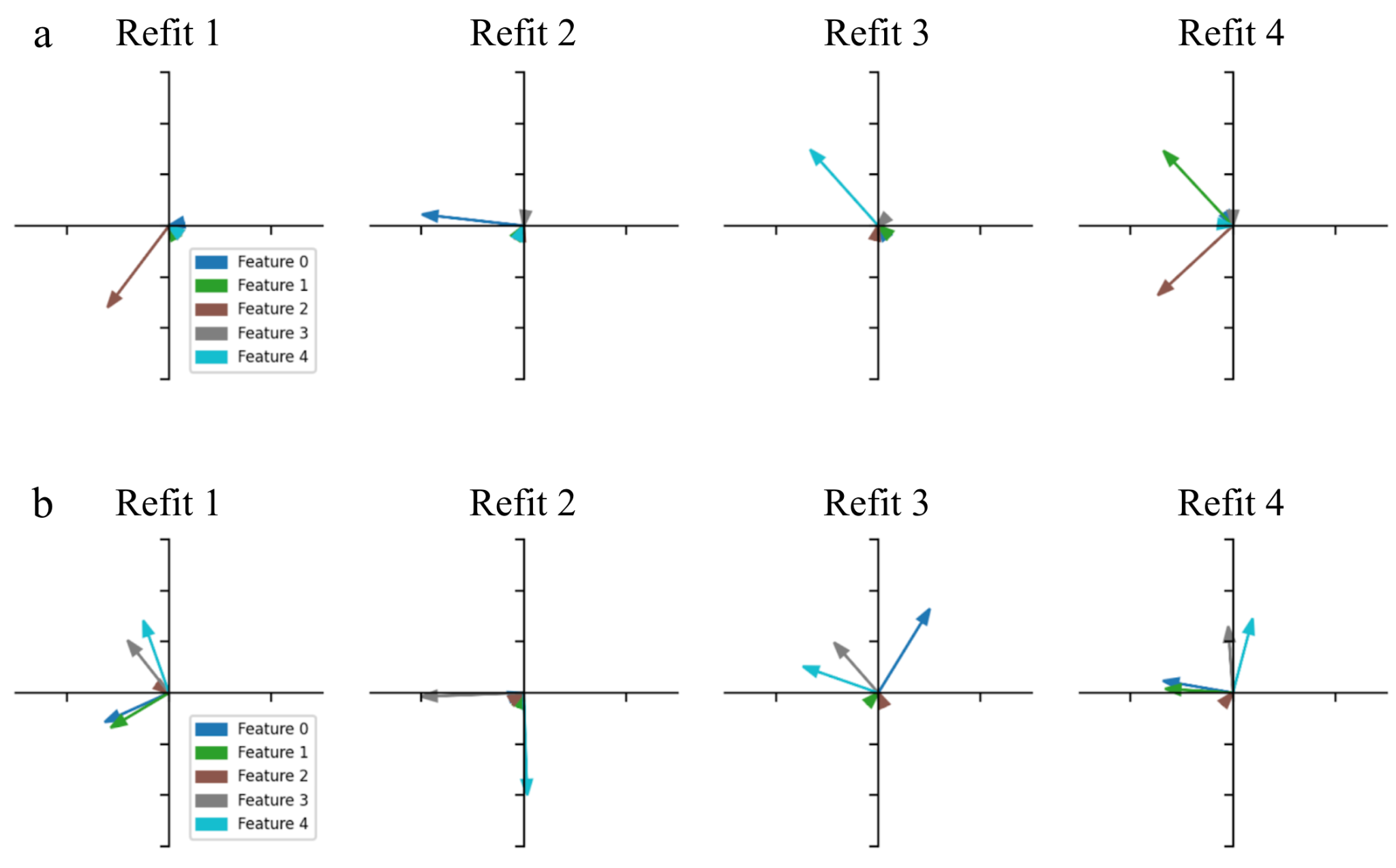

**Supplementary Figure 2. The correlation regime between input features shapes model behavior.** Based on Elhage et al.’s code and findings (Elhage et al. 2022). We train their proposed autoencoder where the latent representation is a linear compression to two dimensions of the inputs x to a lower dimension h = Wx. The decoder then decompresses the input via a ReLU activation x’ = ReLU(W^T^ h + b) = ReLU(W^T^Wx + b). Our dataset is multivariate gaussian where N_features_ = 5 are different components of the gaussian with mean vector zero and unit variance. The covariance between features is controlled via a hyperparameter α, which describes the sparsity of the covariance matrix. The loss function is the mean squared error between inputs and outputs, since we assigned all features equal importance, L = Σ_x_Σ_i_ (x_i_-x_i_’)^2^. Each column in the weight matrix W corresponds to a direction in the latent space (2D) representing the projection of each input feature x_i_ in latent space, which we plot here. **a)** Compressing 5 independent features (α = 0) to 2 latent dimensions, the network learns a maximum of two input features and ignores the rest, mapping them to 0 (the center). Note that the vectors are orthogonal, i.e. 1 dimension is necessary per independent input feature. **b)** Adding correlations between features (α = 0.7), the network can represent more than 2 features, since they are not independent anymore, even when all features are active simultaneously (0 feature sparsity). Here we show four refits of the same model for each of the two correlation regimes.

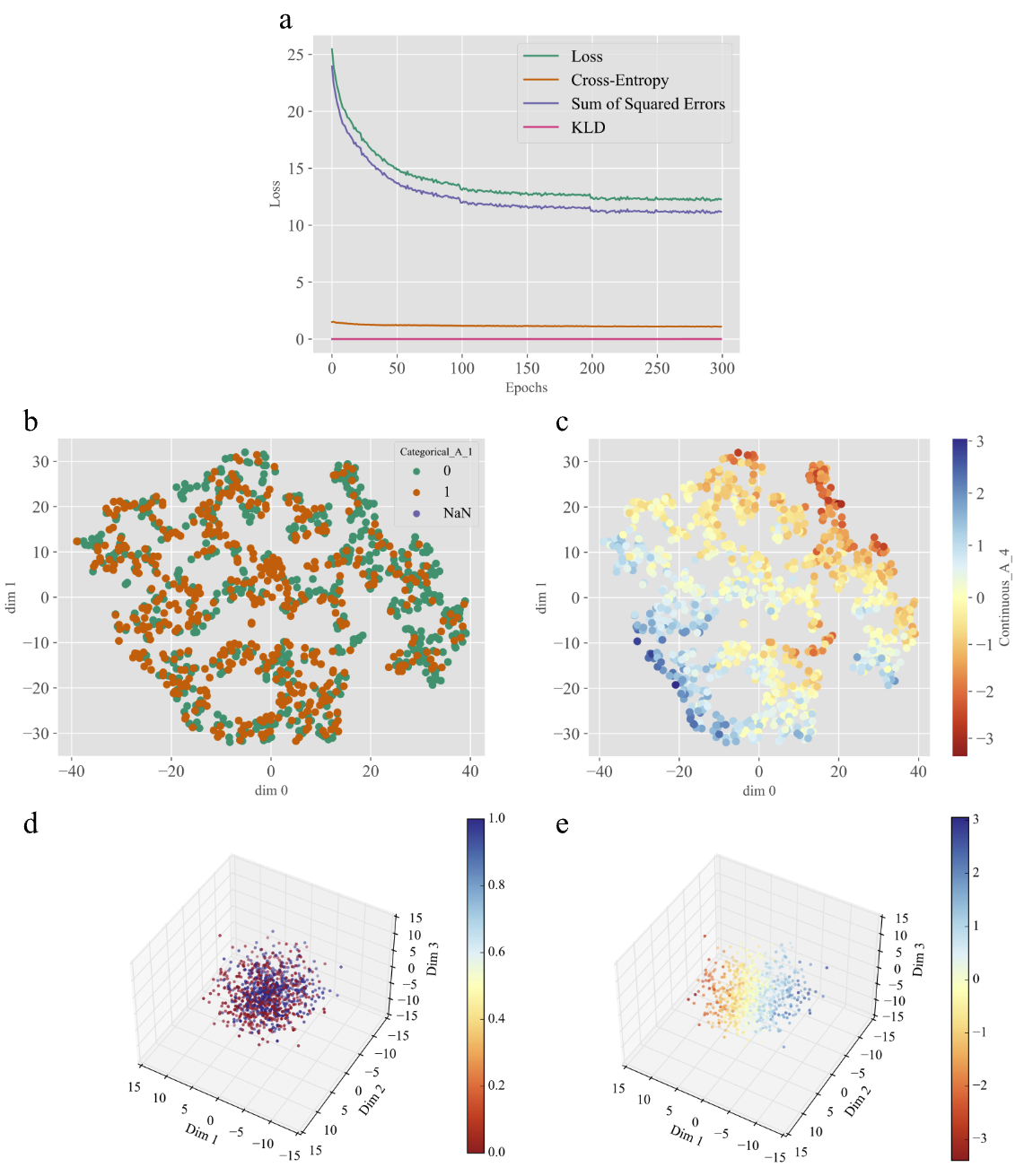

**Supplementary Figure 3.** **The relative importance of each data modality shapes sample organization in latent space.** Upweighting by a factor of 10 the loss for both continuous datasets reduces clustering and promotes a more homogeneous latent space, at the expense of the model ignoring the categorical variables. **a)** Loss curve when upweighting continuous dataset contributions. Categorical loss, in orange, remains almost constant, illustrating that the model does not learn how to encode and reconstruct the categorical data. Loss landscape and optimization is hence shaped by the continuous variables. **b, c)** Umap representations of the latent space color coded by the labels for b) Categorical_A_1 and c) Continuous_A_4. **d, e)** Complete representation of the 3D latent space, color coded according to the same features as in b) and c). Note that the latent space is homogeneously populated, samples are ordered following the value gradient of the learned continuous feature, but there is no easy boundary separating categories which complicates their reconstruction. Note also that the KLD term is zero, not due to a posterior collapse, but instead because it is weighted by β and therefore does not contribute to the overall loss.

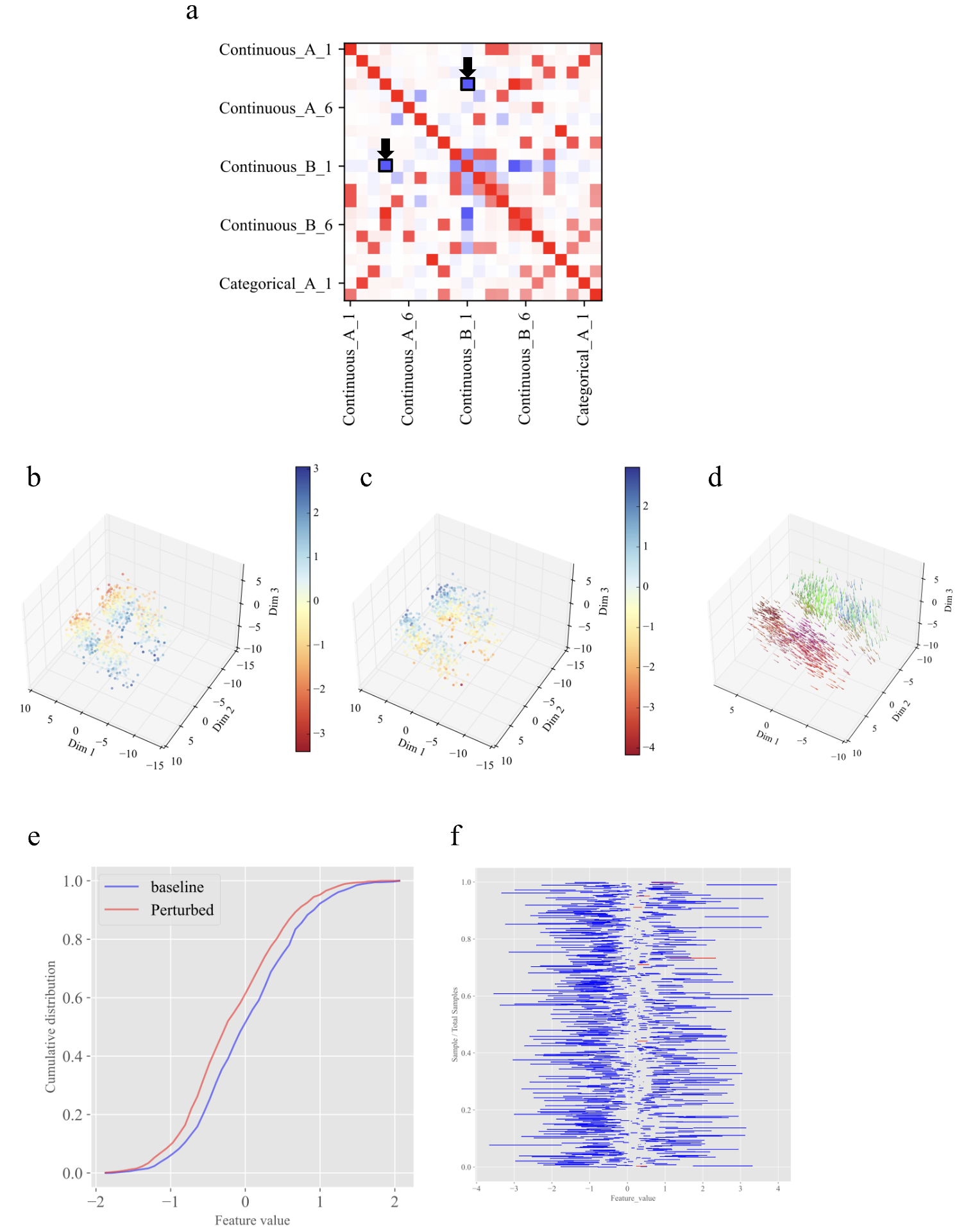

**Supplementary Figure 4. Effects of perturbing associated input variables. a)** Ground truth associations between features for the dataset with added correlations (III). Continuous_A_4 and Continuous_B_1 display a strong inverse correlation. **b, c)** Latent space sample representations color coded by Continuous_A_4 and Continuous_B_1 values respectively. Samples/regions for which A_4 is high are also samples/regions for which B_1 is low and vice versa. **d)** Perturbing positively Continuous_A_4 brings samples with low Continuous_A_4 (high Continuous_B_1) towards the region where we have samples with high Continuous_A_4 (low Continuous_B_1). **e)** Cumulative distributions of the reconstructed values for the feature Continuous_B_1 across samples when feeding the original samples to the encoder (baseline) and when using as inputs the samples where Continuous_A_4 has been perturbed positively. One can notice that the cumulative distribution of reconstructed feature values presents a negative shift (KS distance = 0.134) **f)** Plot showing the increase or decrease in value for the feature of interest (Continuous_B_1) in all samples (y axis) when perturbing another feature, in this case Continuous_A_4. Red lines show increases in value (left to right) while blue lines indicate decreases (right to left). Perturbing a feature associated with the one measured causes most of the samples to replicate the direction of change with varying magnitudes.

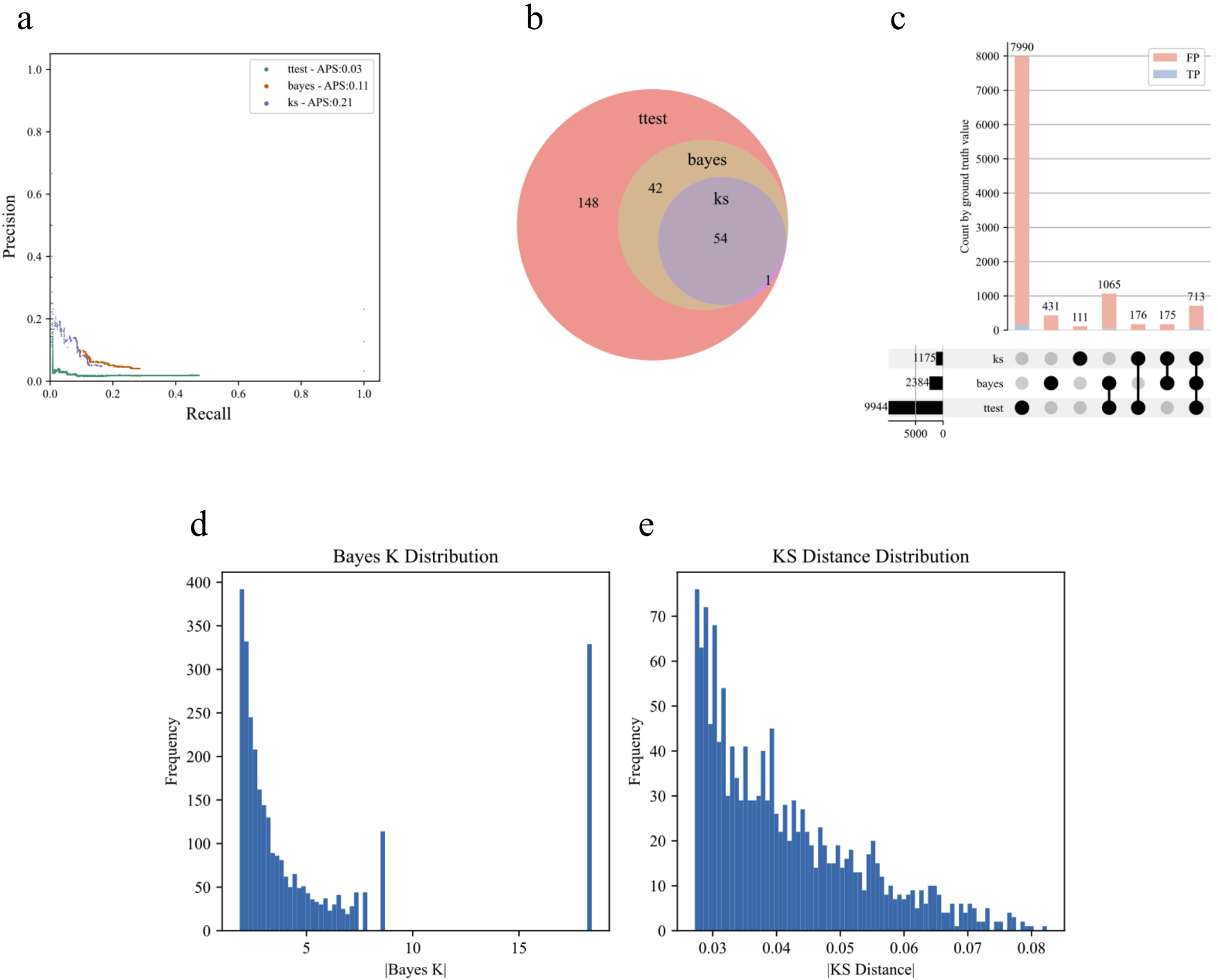

**Supplementary Figure 5. Performance comparison between methods. a)** Precision recall curves for the different methods. **b)** Overlap between true positive features found by the different models. **c)** UpSet plot of all associations found by the different methods and fraction of correct associations. Note that the t-test method has a high false positive rate. **d)** Bayes score value distribution. Bayes_K = 18.42 is obtained when all samples in the cohort increase or decrease in value. **e)** KS score distribution.

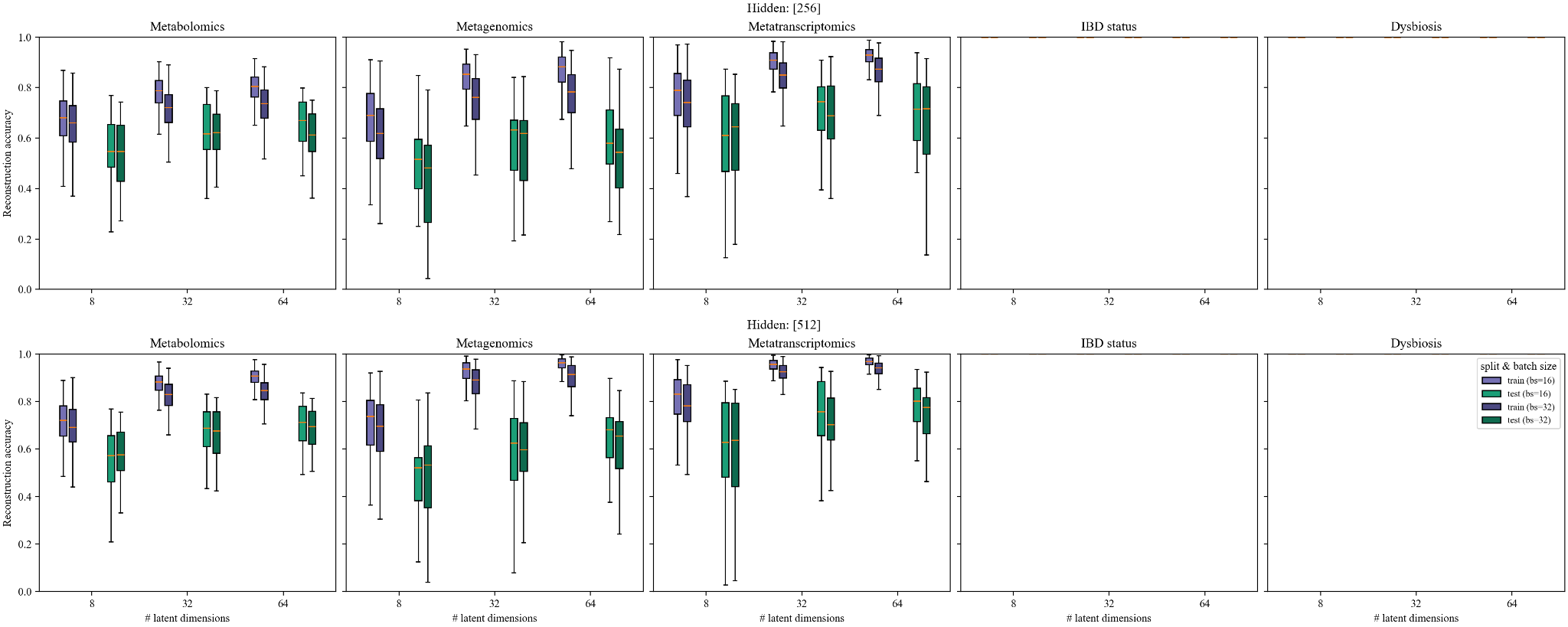

**Supplementary Figure 6. Hyperparameter tunning.** We fixed the number of epochs to 400 and β=1e-4 to ease the reconstruction. We explored combinations of batch size={16,32}, n_hidden_nodes ={256,512} and n_latent_nodes = {8,32,64}. Note that both categorical variables (IBD status and Dysbiosis) were perfectly reconstructed regardless of the hyperparameters (flat orange lines at the top border of the plots).

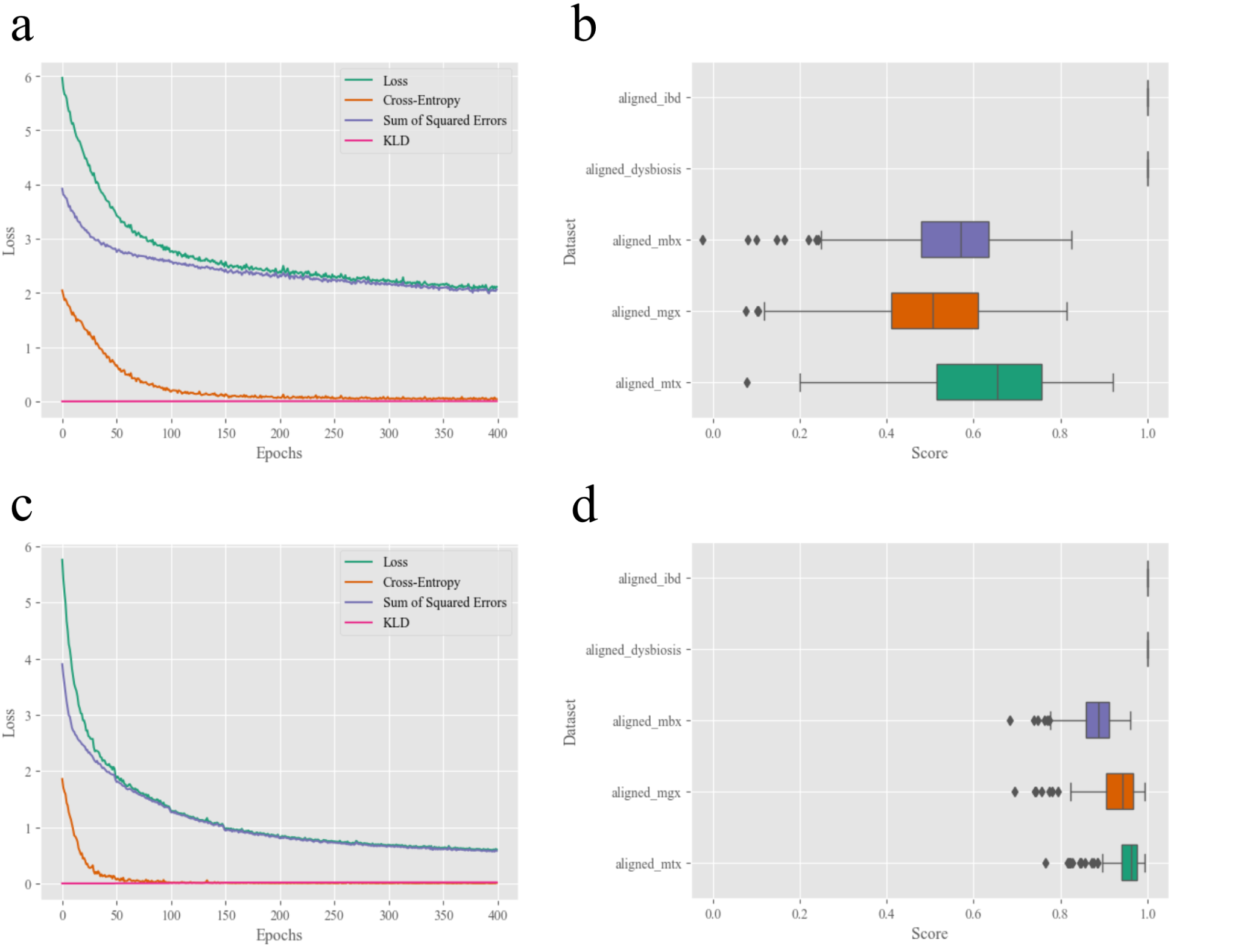

**Supplementary Figure 7. Loss and reconstruction accuracy. a,b)** Results for a MOVE model with n_latent_ = 16, n_hidden_ = 64 and β = 0.0001. **a)** Loss evolution decomposed as a continuous contribution (Sum of Squared Errors), categorical contribution (Cross-Entropy) and a negligible regularization on the latent space distribution (KLD). **b)** Reconstruction accuracy given by cosine similarity. **c,d)** Results for a MOVE model with n_latent_ = 32, n_hidden_ = 512 and β =0.0001. Note that the categorical loss goes to zero first, i.e. the model perfectly reconstructs categorical variables by ordering the latent space in easily separable clusters. Also note that the KLD term is zero not due to a posterior collapse, but instead because it is weighted by β and therefore does not contribute to the overall loss.

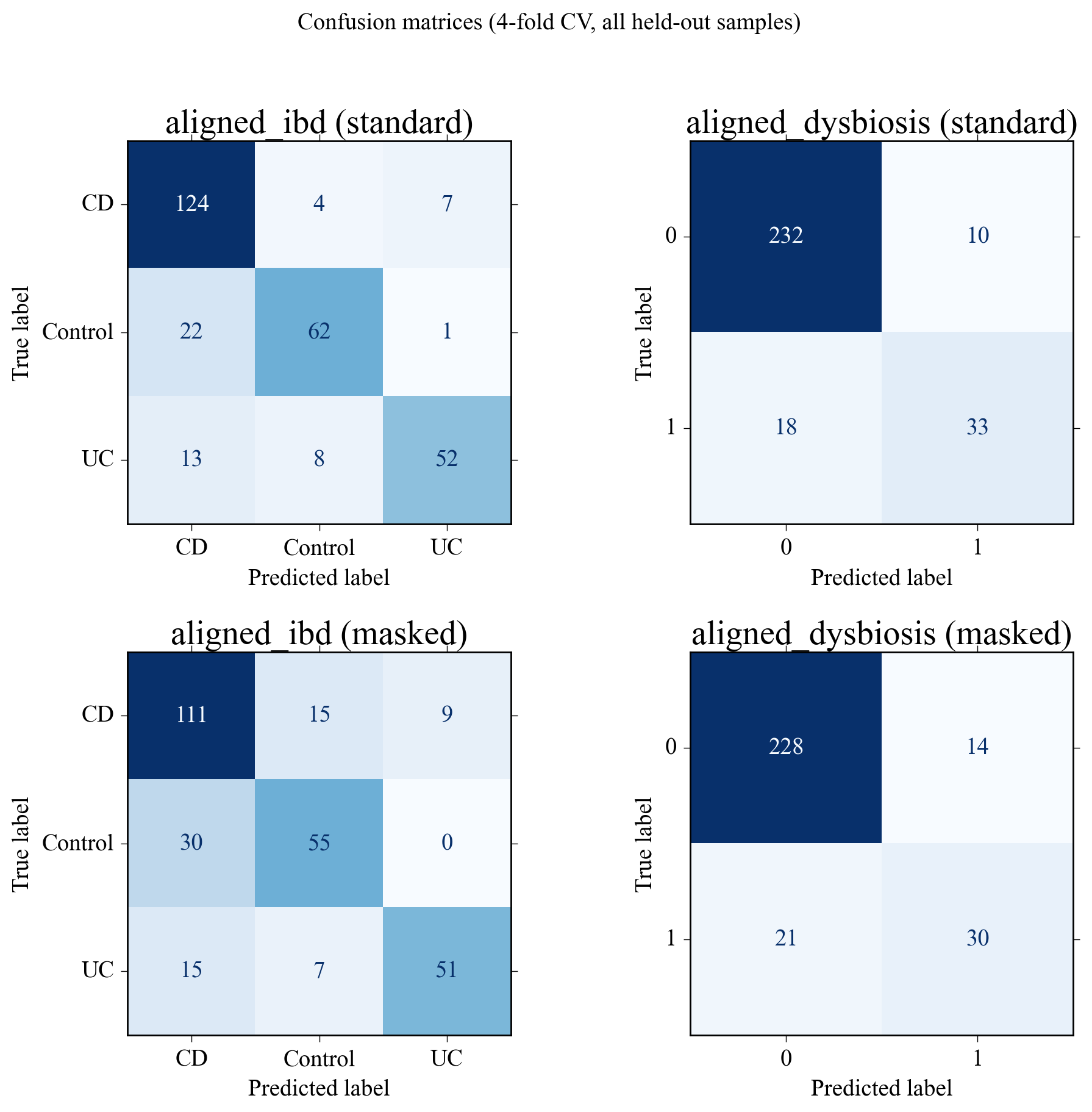

**Supplementary Figure 8. Confusion matrices for the held-out sample predictions in the IBDMDB dataset in a 4-fold cross validation analysis.** This plot illustrates the role of the decoder as an implicit classifier of the latent representations of the samples towards their categorical labels. **Top row)** Decoder classification accuracy when categorical variables were still used as inputs to the encoder (standard). **Bottom row)** Decoder classification accuracy when masking the categorical variables from the inputs to the encoder (masked). Performance drops slightly, but that the rest of the variables (continuous) still carry enough information to make the classification possible. Please note that this behavior is specific to this dataset, i.e. continuous variables do not necessarily encode information about the categorical labels in other datasets.

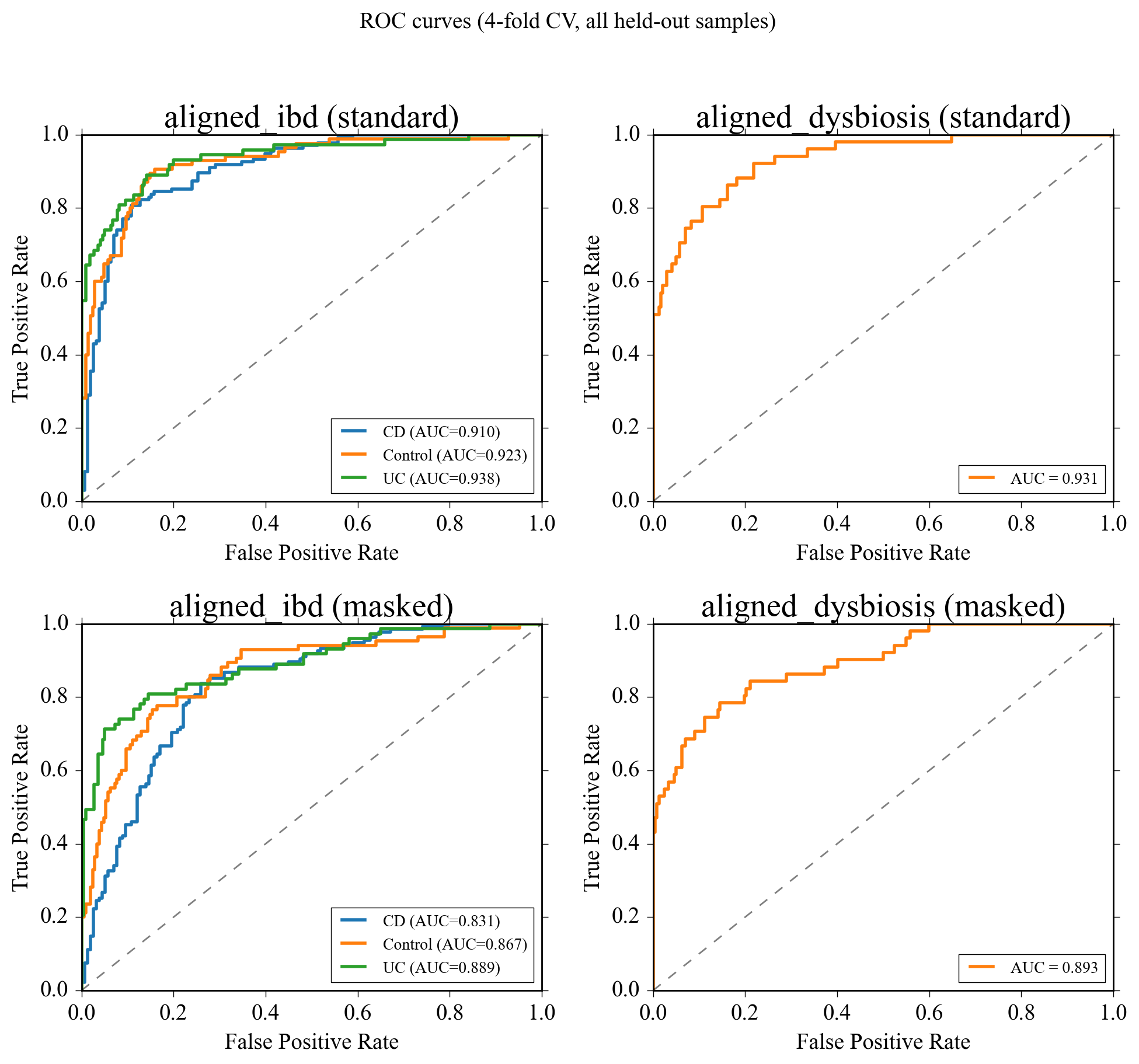

**Supplementary Figure 9. ROC curves when predicting IBDMDB labels using MOVE. Top row)** Providing (standard) and **Bottom row)** masking (masked) the categorical labels to MOVE’s encoder when predicting IBDMDB labels. Please note that the classes are highly imbalanced.

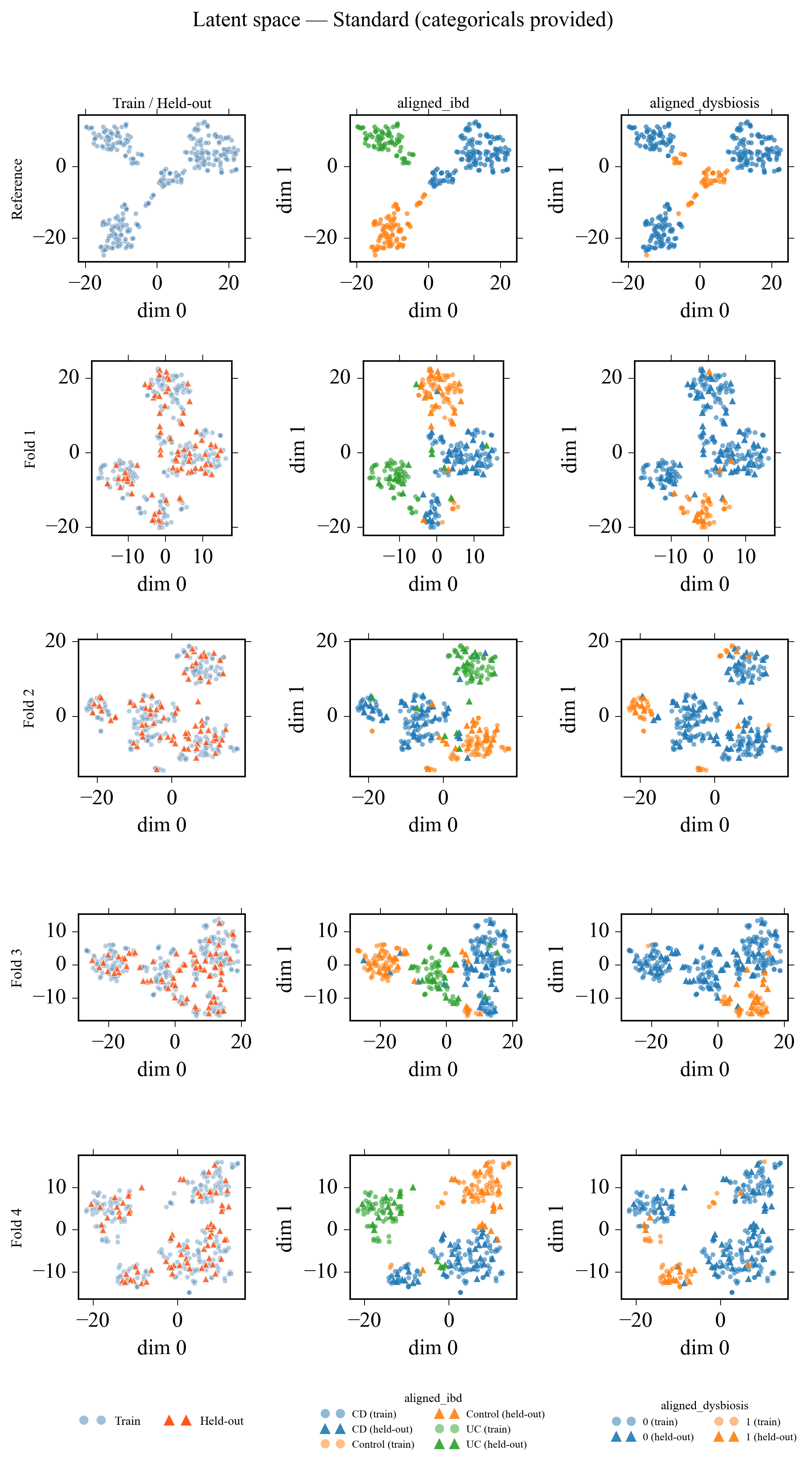

**Supplementary figure 10. Latent space representations of the IBDMDB samples when categorical variables are provided as inputs to the model.** Providing the categorical labels to the model and asking it to properly reconstruct them incentivizes the encoder to separate samples corresponding to different classes into well-defined clusters.

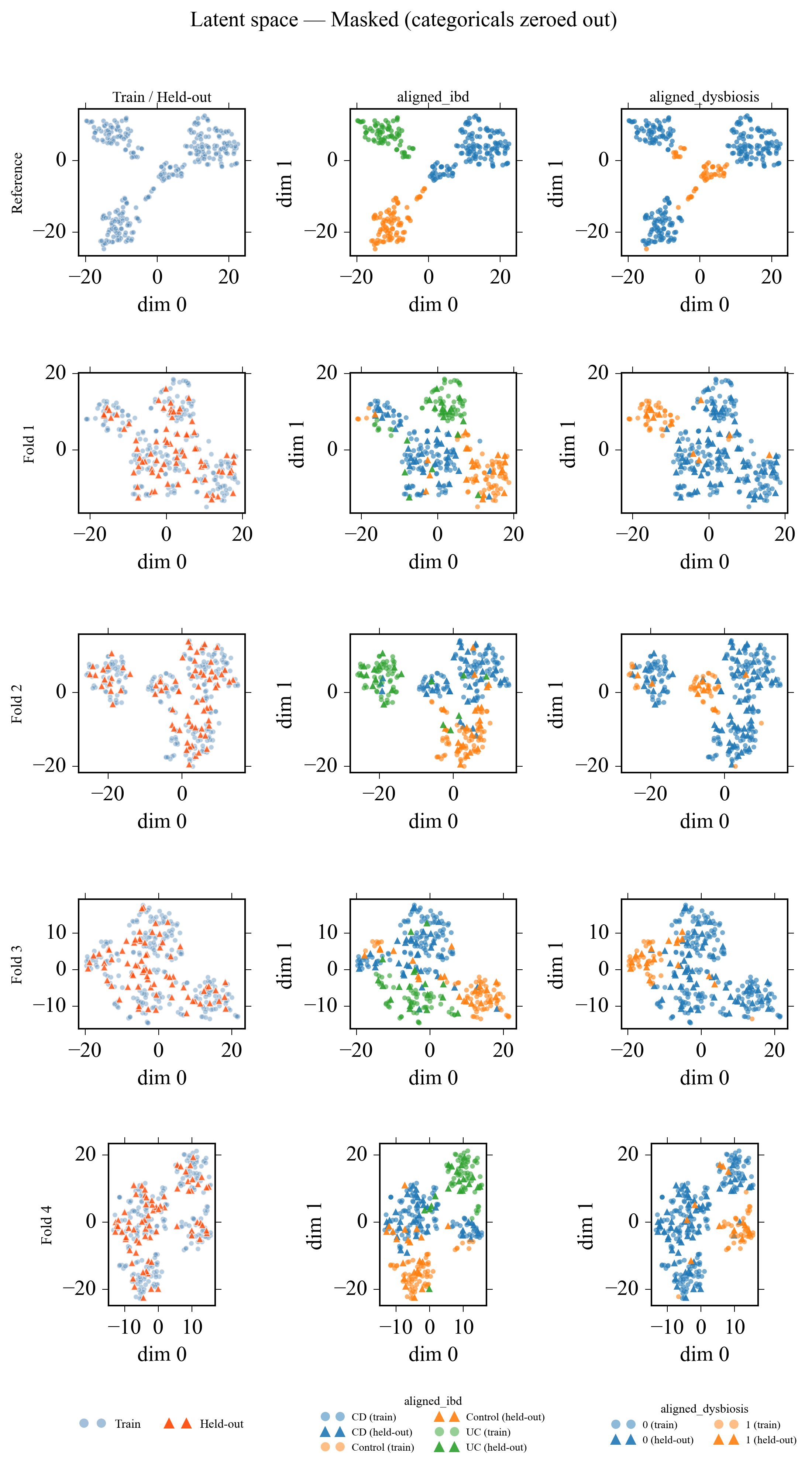

**Supplementary figure 11. Latent space representations of the IBDMDB samples when categorical variables are masked, i.e. not provided explicitly to the model.** Masking categorical features leads to a less pronounced clustering and hence a slight drop in the decoder’s classification power (Figure Conf Mat). For this particular dataset, the rst of the variables still capture enough information about the classes to be able to map the samples to the right region and hence enable the downstream classification by the decoder.

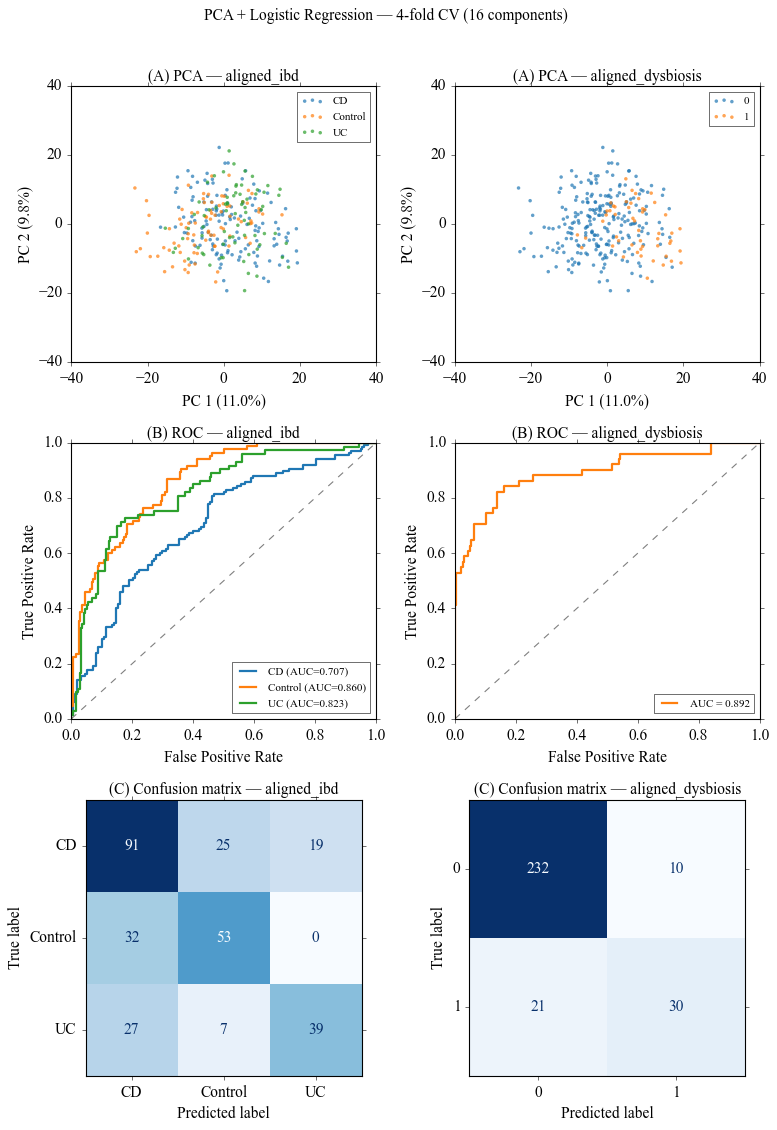

**Supplementary Figure 12. PCA analysis on concatenated continuous profiles and subsequent logistic regression to disease status and dysbiosis in the IBDMDB dataset.** ROC curves and confusion matrices were computed on held out samples in the same 4-fold cross-validation scheme. **Top row)** Sample profiles containing all continuous modalities (mgx, mtx and mbx) were compressed to 16 components to match MOVE’s latent dimensionality. **Middle row)** Receiver Operator Characteristic (ROC) curves. **Bottom row**) Confusion matrices. Note that, for this dataset, the continuous profiles are highly predictive of the corresponding categorical labels.

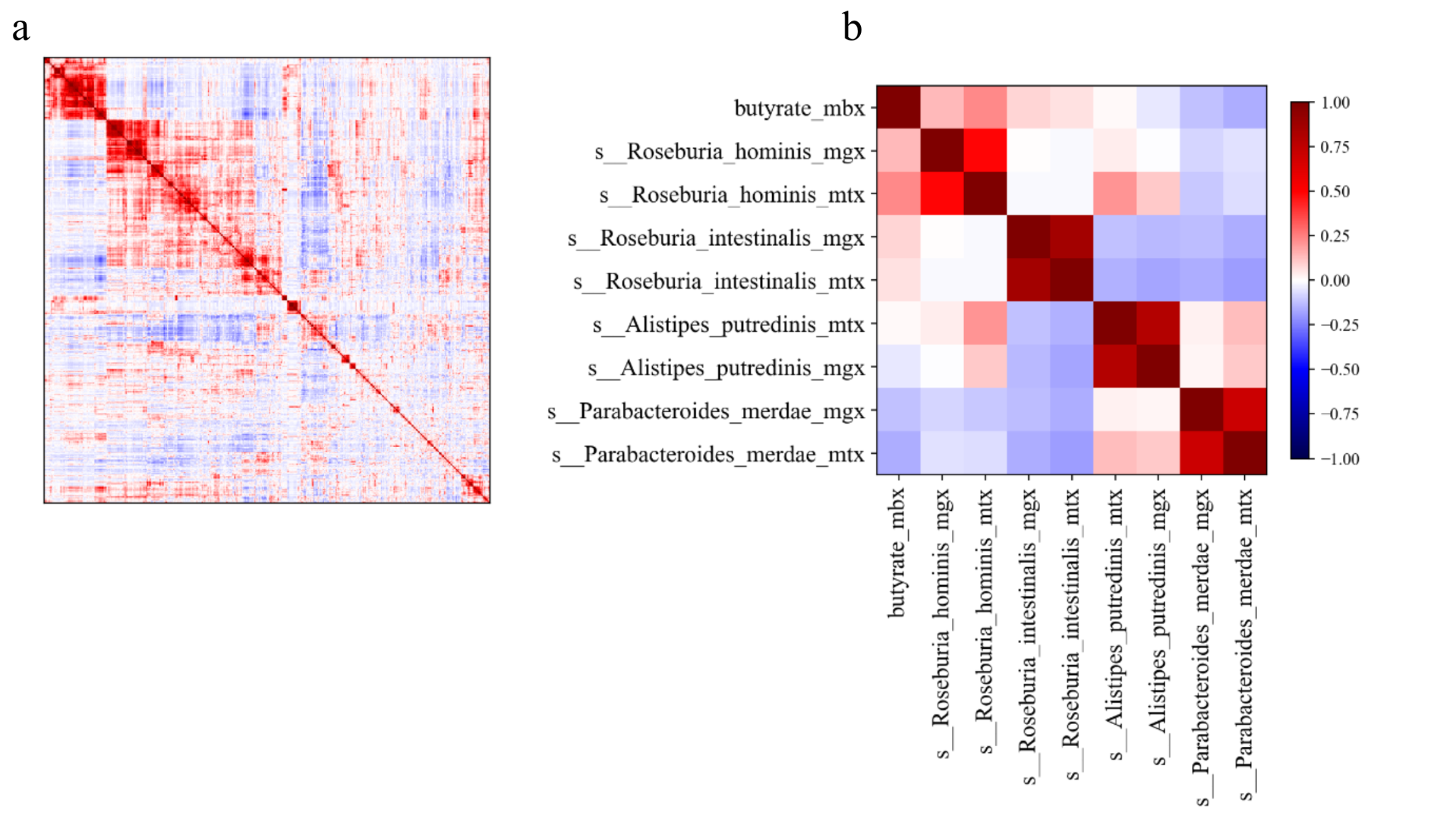

**Supplementary Figure 13**. **Correlational structure in the data.** **a)** Cross-modality correlations between metabolomics, metatranscriptomics and metagenomics in the IBDMDB data after filtering. **b)** Cross-modality correlations for the same species. Please note the strong coupling between metagenomics and metatranscriptomics profiles for the same species, but also the close similarity between metabolite profiles and the bacterial species that produce them, e.g. *R. hominis* and butyrate (Lloyd-Price et al. 2019).

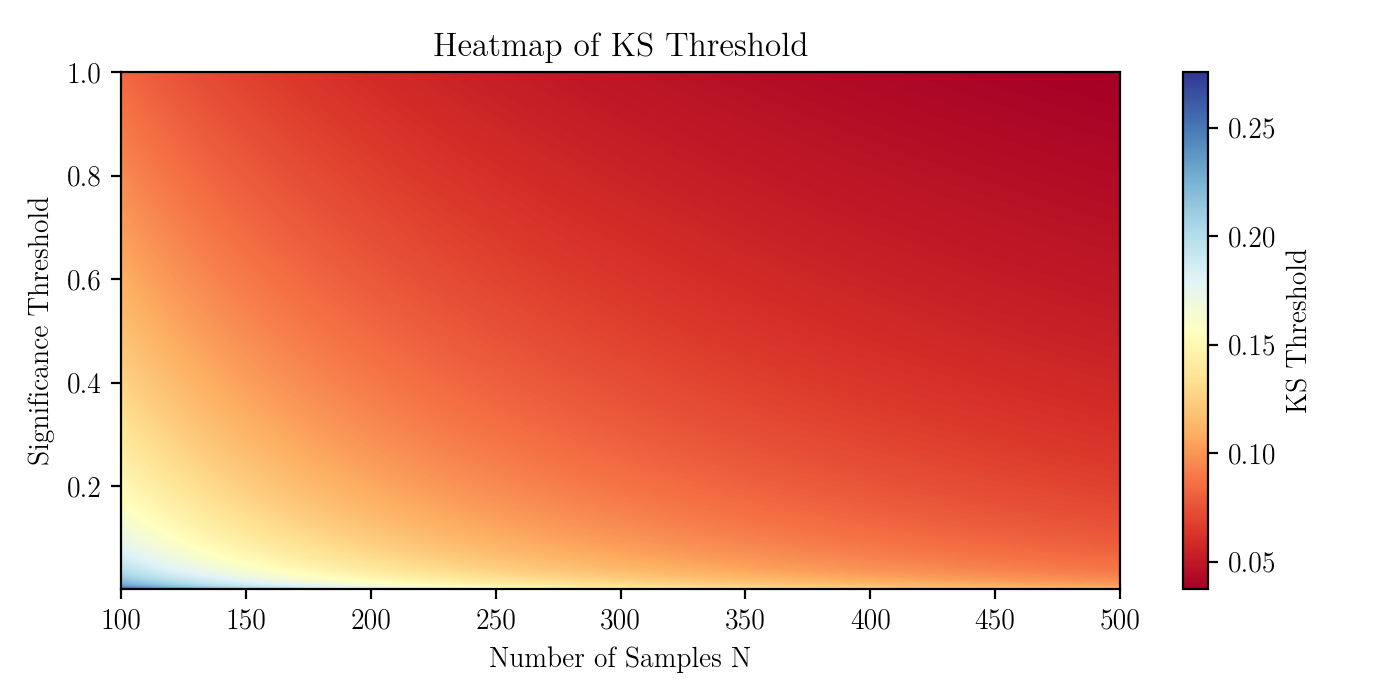

**Supplementary Figure 14. KS threshold strongly depends on the number of samples available.** Heatmap showing the KS distance threshold ($D_{n}$) from which the reconstructed distributions before and after a perturbation of a certain feature can be considered to be different, as a function of certain confidence (Significant threshold $\alpha$) and the number of available samples ($n$) $D_{n}> \sqrt{-\frac{1}{n} \cdot ln\left( \frac{\alpha}{2} \right)}$. In most cases, the effect of perturbing a feature in the input does not alter the reconstructed distribution enough to consider them different distributions, especially under a low sampling regime.

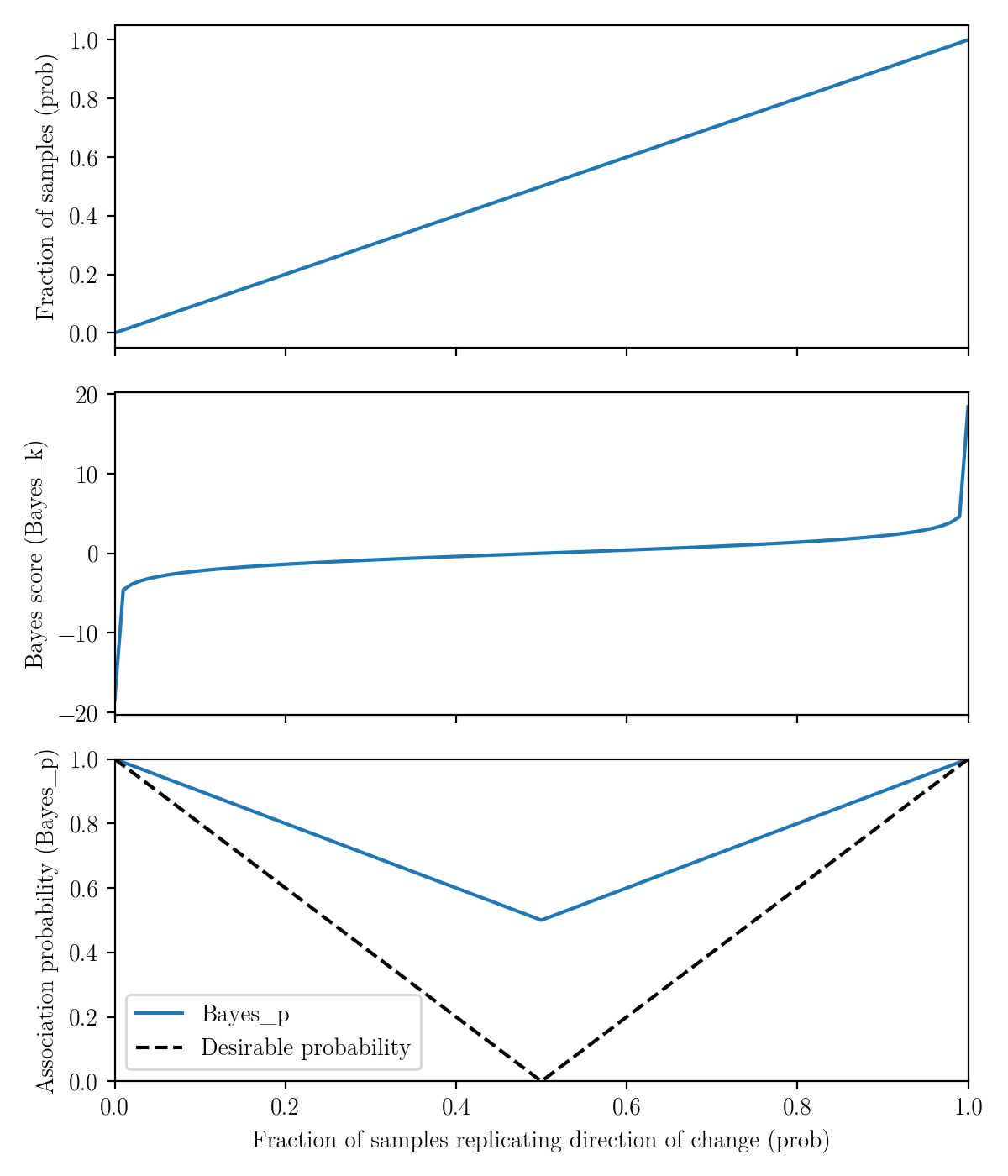

**Supplementary Figure 15. Bayesian association criteria reduce significance barriers for perturbation effect detection.** The Bayes-inspired approach quantifies associations between perturbed feature A and target feature B based on the fraction of samples where B increases upon a perturbation in A (prob, x-axis). The plots show the theoretical link between any sample fraction reproducing a change (prob), the corresponding Bayes score (Bayes_k) and the matching association probability (Bayes_p). Note that associations can reach 100% probability for minimal changes across all samples if they all change in one direction, making the method more suitable to quantify subtle reconstruction changes. **Top)** The contribution of highly confident associations (prob={0,1}) to the cumulative evidence is driven by the stability constant ε, which induces that sharp change of Bayes_k for prob=0 and prob=1 (**Methods**) **Bottom)** The final Bayes_p value is interpreted as the probability of A-B association after applying the transformations presented in Methods and Allesøe et al. (Allesøe et al. 2023). Note that all possible feature pairs have a minimum association probability Bayes_p = 0.5 instead of a desirable 0, with prob=0.5 indicating random B changes upon A perturbation.

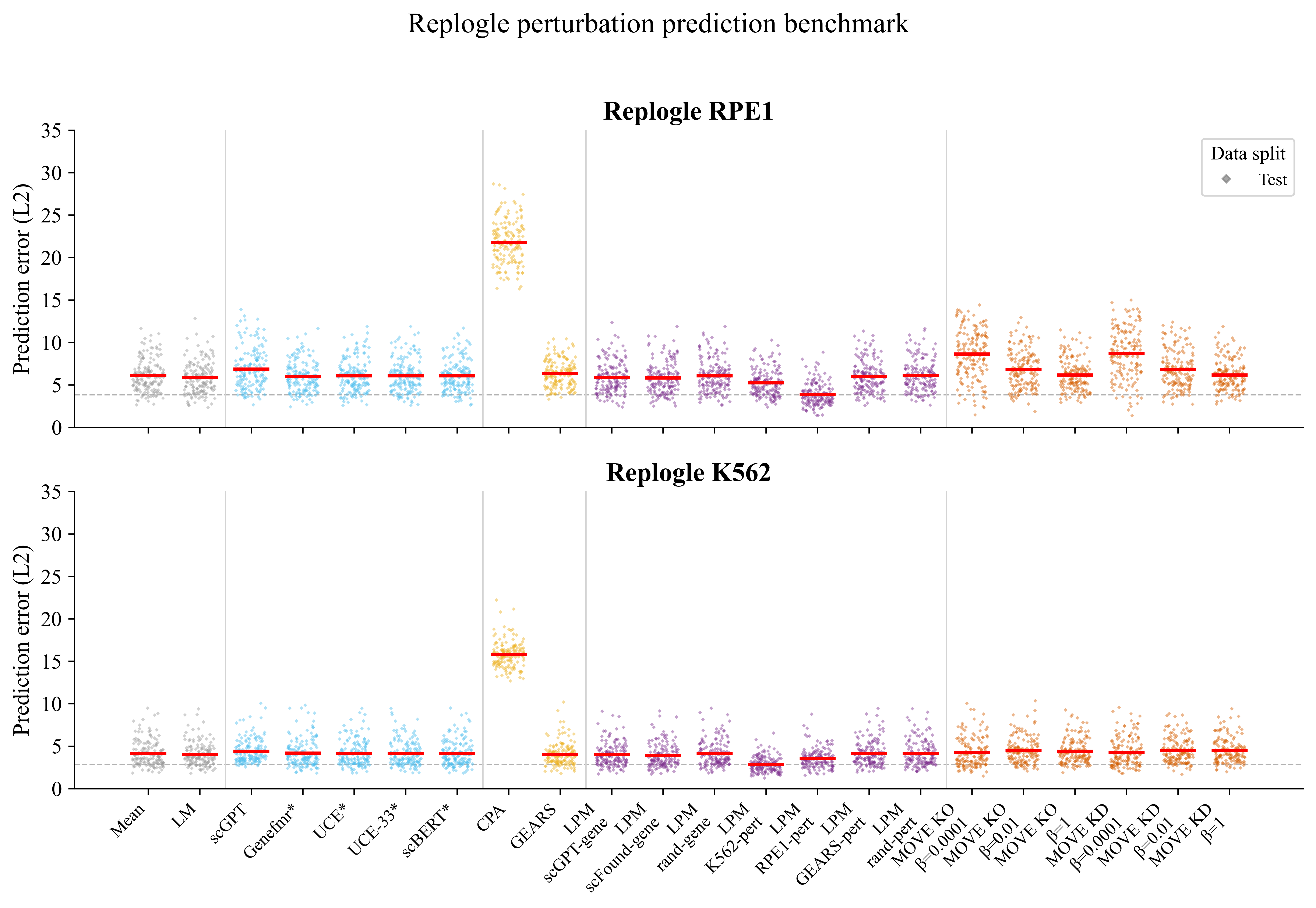

**Supplementary Figure 16. L2 score for each single KD experiment in the test set.** The same gene KDs are shown across all models. Benchmark data was obtained from the source data file for single gene perturbations from Ahlmann-Eltze et al., Panel C. The MOVE model shown here is the same as in the main figures, with 64 hidden nodes and 16 latent nodes. Different regularization regimes ($\beta$= {0.0001,0.01,1}) and perturbation schemes (KD $\to$ minus std, KO $\to$ minimum across training samples) are shown. The L2 distance for MOVE predictions is comparable to that of the other approaches.

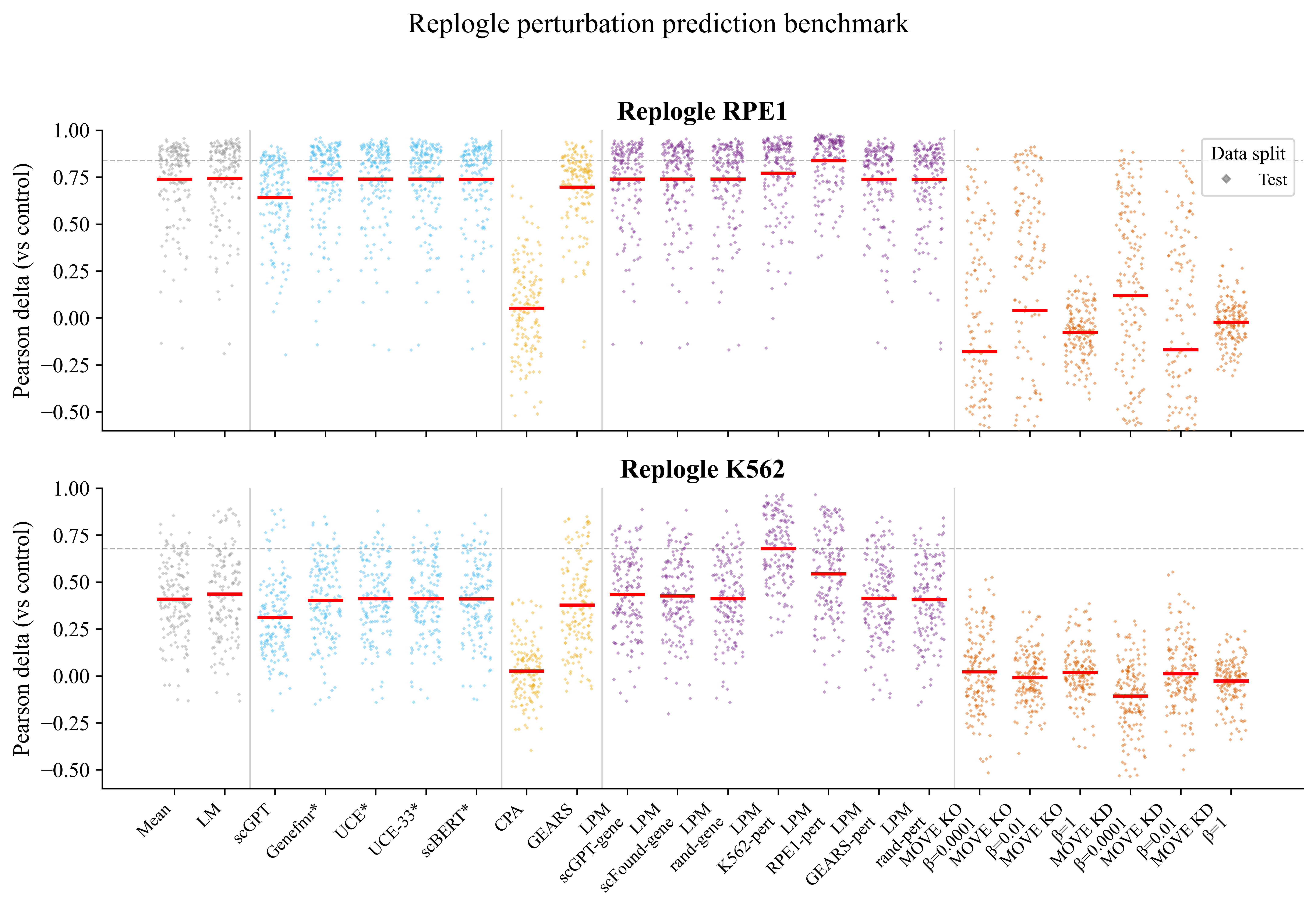

**Supplementary Figure 17. Pearson delta compared with controls for each single KD experiment in the test set.** The same gene KDs are shown across all models. Benchmark data for the other models was found in the source data file describing single gene perturbations from Ahlmann-Eltze et al., Panel C. The MOVE model shown here is the same as in the main figures, with 64 hidden nodes and 16 latent nodes. Different regularization regimes ($\beta$= {0.0001,0.01,1}) and perturbation schemes (KD $\to$ minus std, KO $\to$ minimum across training samples) are shown. VAE-based approaches underperformed the linear baselines both when introducing perturbations in input space (MOVE) or in latent space (CPA).

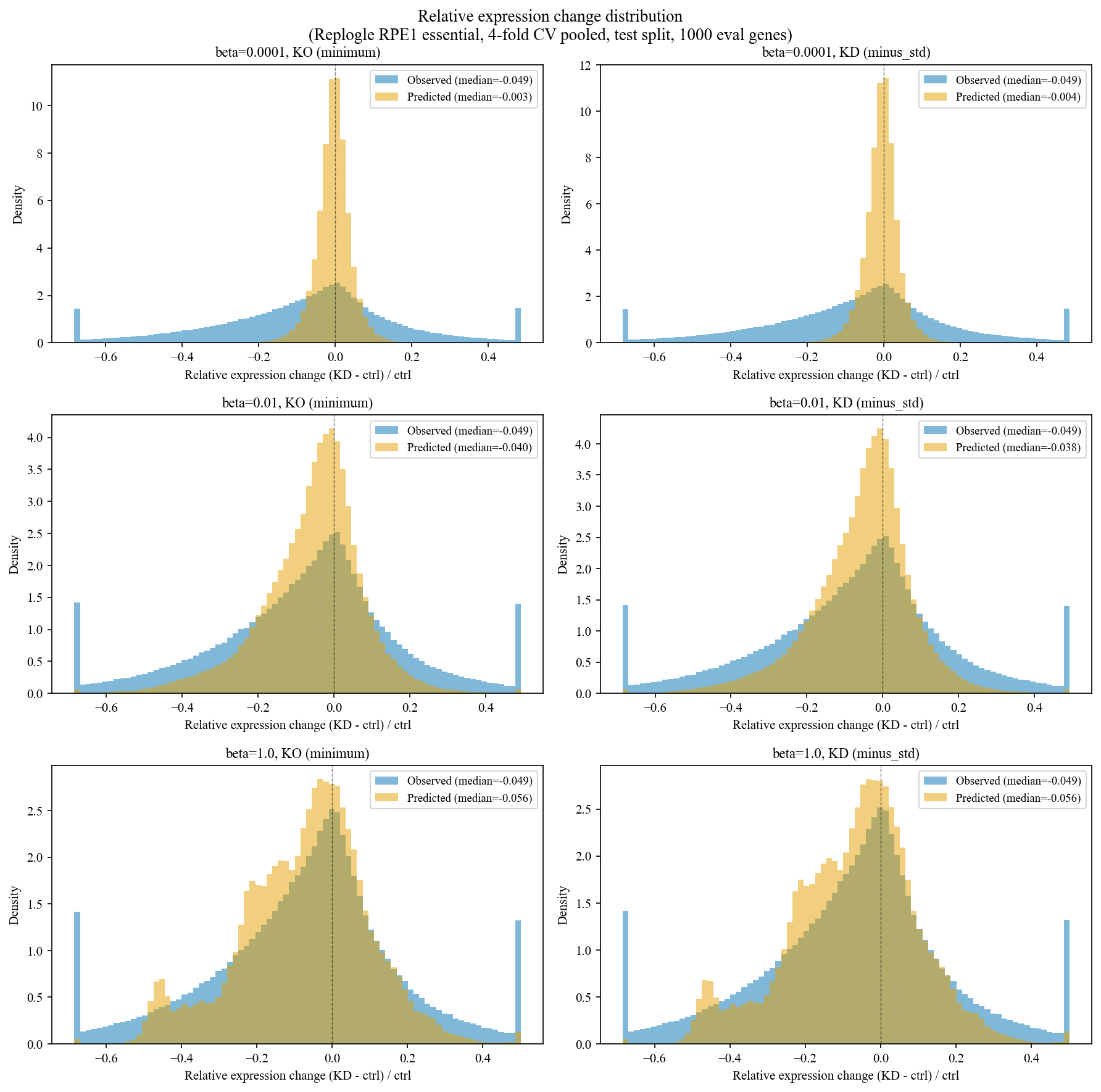

**Supplementary Figure 18. Predicted and Observed expression changes in Replogle RPE1 essential dataset (obtained from GEARS and untransformed).** Test split predictions in a 4-fold cross validation, measuring 1000 evaluation genes. The magnitude of the predicted changes is smaller than the magnitude of real changes, seen as a narrower distribution. This effect is more pronounced at low regularization regimes. Expression changes after a KD were more pronounced in RPE1 cells than in K562 cells.

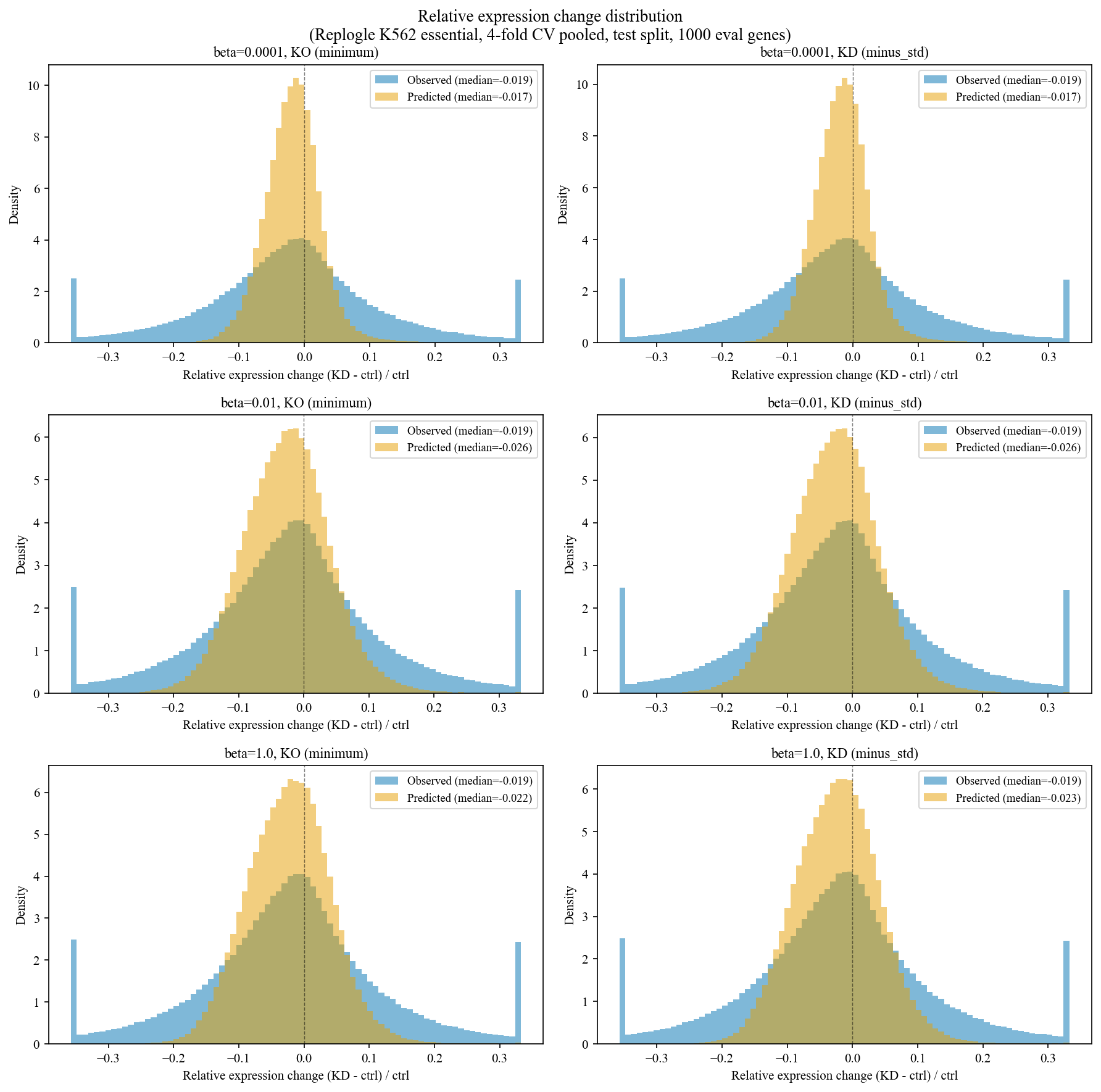

**Supplementary Figure 19. Predicted and Observed expression changes in Replogle K562 essential dataset (obtained from GEARS and untransformed).** Test split predictions in a 4-fold cross validation, measuring 1000 evaluation genes. The magnitude of the predicted changes is smaller than the magnitude of real changes, seen as a narrower distribution. This effect is more pronounced at low regularization regimes. Expression changes after a KD were more pronounced in RPE1 cells than in K562 cells.

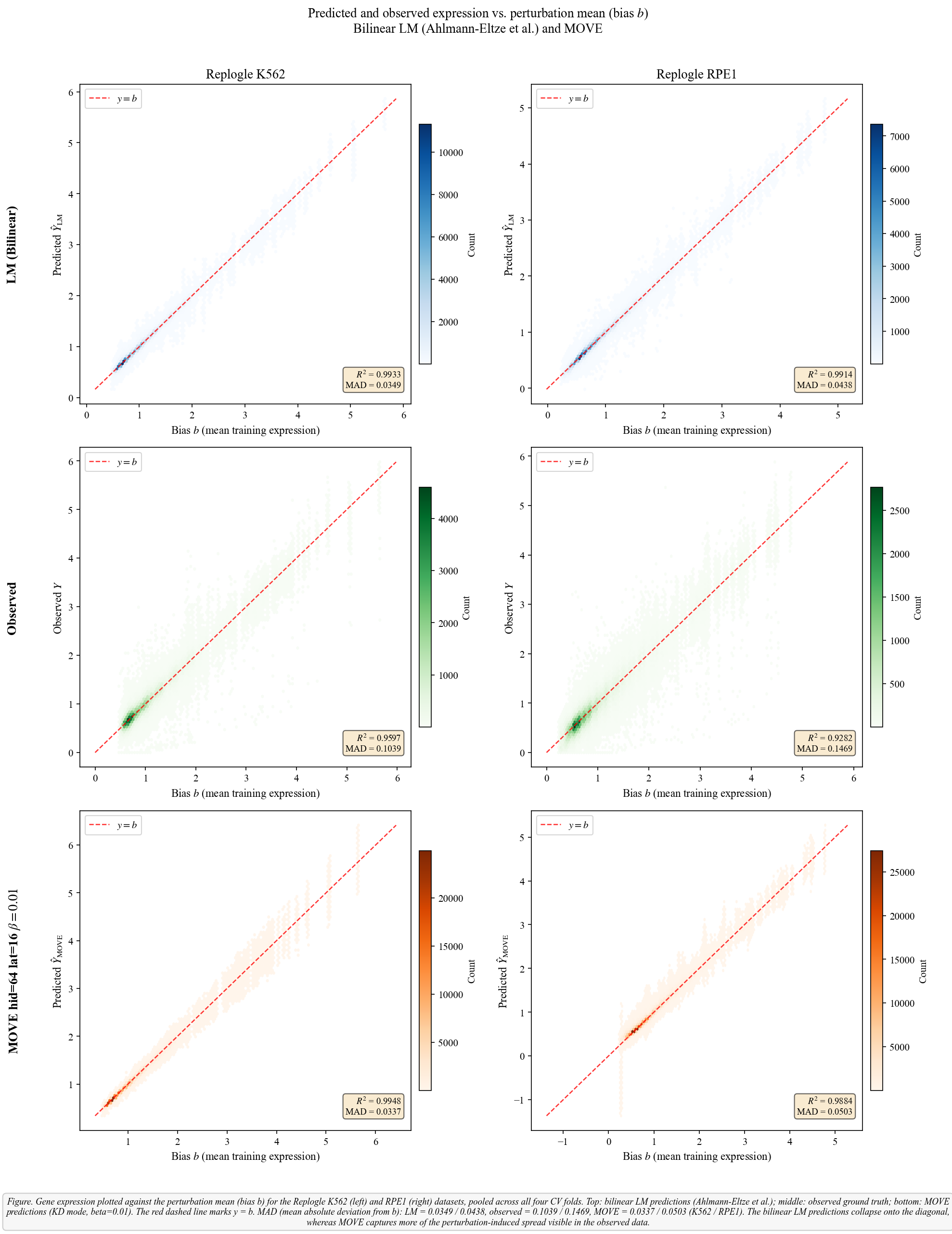

**Supplementary Figure 20. Gene expression plotted against the bias term b (mean expression across training perturbations) for the Replogle K562 (left) and RPE1 (right) datasets.** Each panel shows results pooled across all four CV folds, the 1000 genes with highest expression in control cells are used for the evaluation, and we show the KD genes covered within the 1000 evaluation genes. The red dashed line marks y = b**. Top row (blue)**: predictions from Ahlmann-Eltze's bilinear model (Y = GWP^T^ + b); **middle row (green)**: observed ground truth; **bottom row (orange)**: MOVE predictions (KD mode, beta = 0.01, 16 latent nodes, 64 hidden nodes). MAD (mean absolute deviation from b) quantifies how far values deviate from the bias. MAD is 0.0349 (K562) and 0.0438 (RPE1) confirming that the bilinear model addition (GWP^T^) barely shifts expression away from the training mean (b). The observed ground truth shows larger MAD values of 0.1039 (K562) and 0.1469 (RPE1) reflecting real perturbation effects that the linear model does not capture. MOVE predicted perturbation effects are also minimal with MAD values of 0.0337 (K562) and 0.0503 (RPE1).

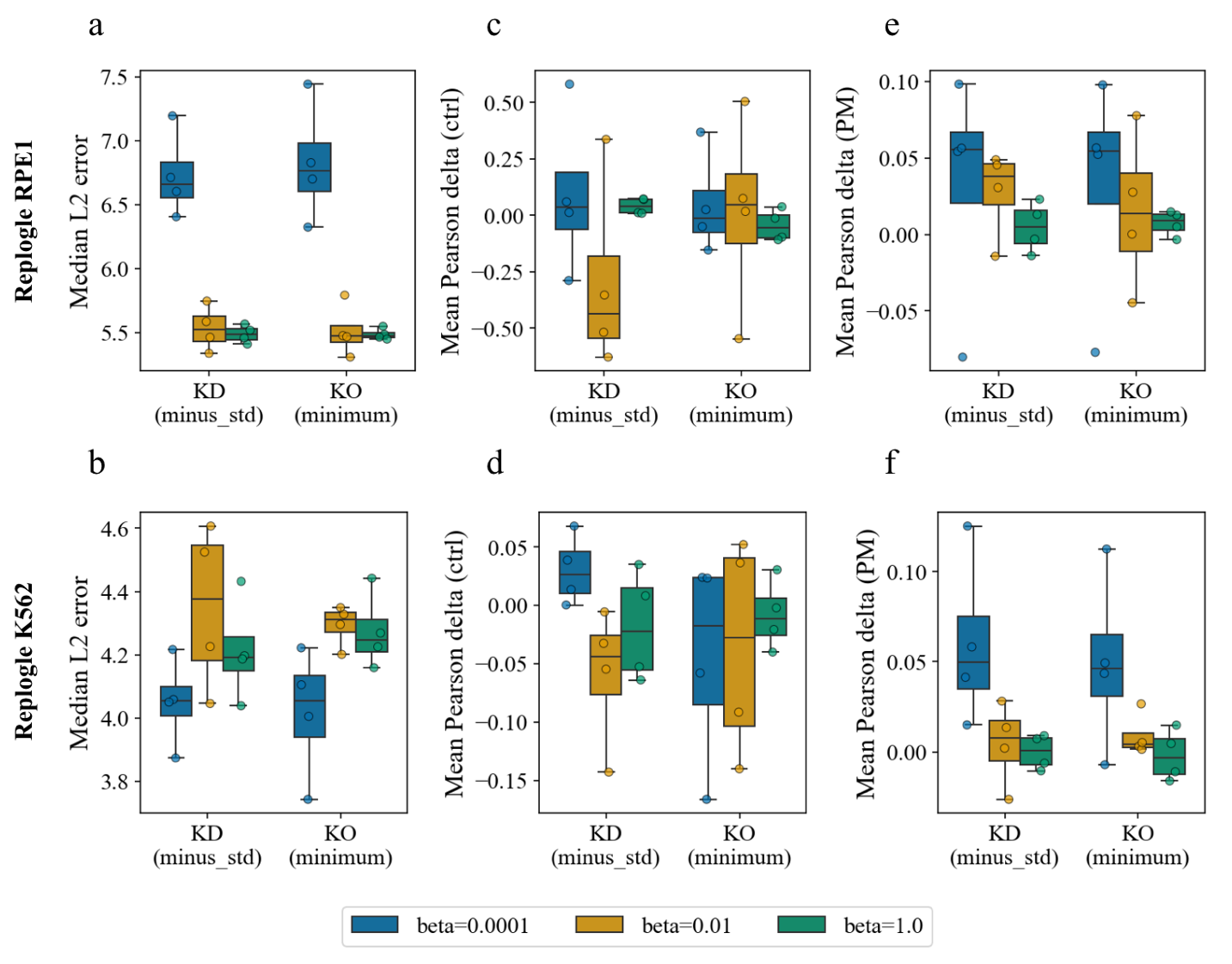

**Supplementary Figure 21. Evaluation of *in silico* input perturbations on single cell CRISPR KD data, different run of the same model architecture (hid=64, lat = 16. a,b.** Median L2 test error across 4 folds under different regularization schemes. **c, d.** Mean Pearson delta measured as the correlation between predictions and observations after subtracting the expression in the control condition. **e, f.** Mean Pearson delta measured as the correlation between predictions and observations after subtracting the mean expression profile across perturbations. L2 and Pearson delta compared to perturbation mean are more reproducible than delta vs. control.

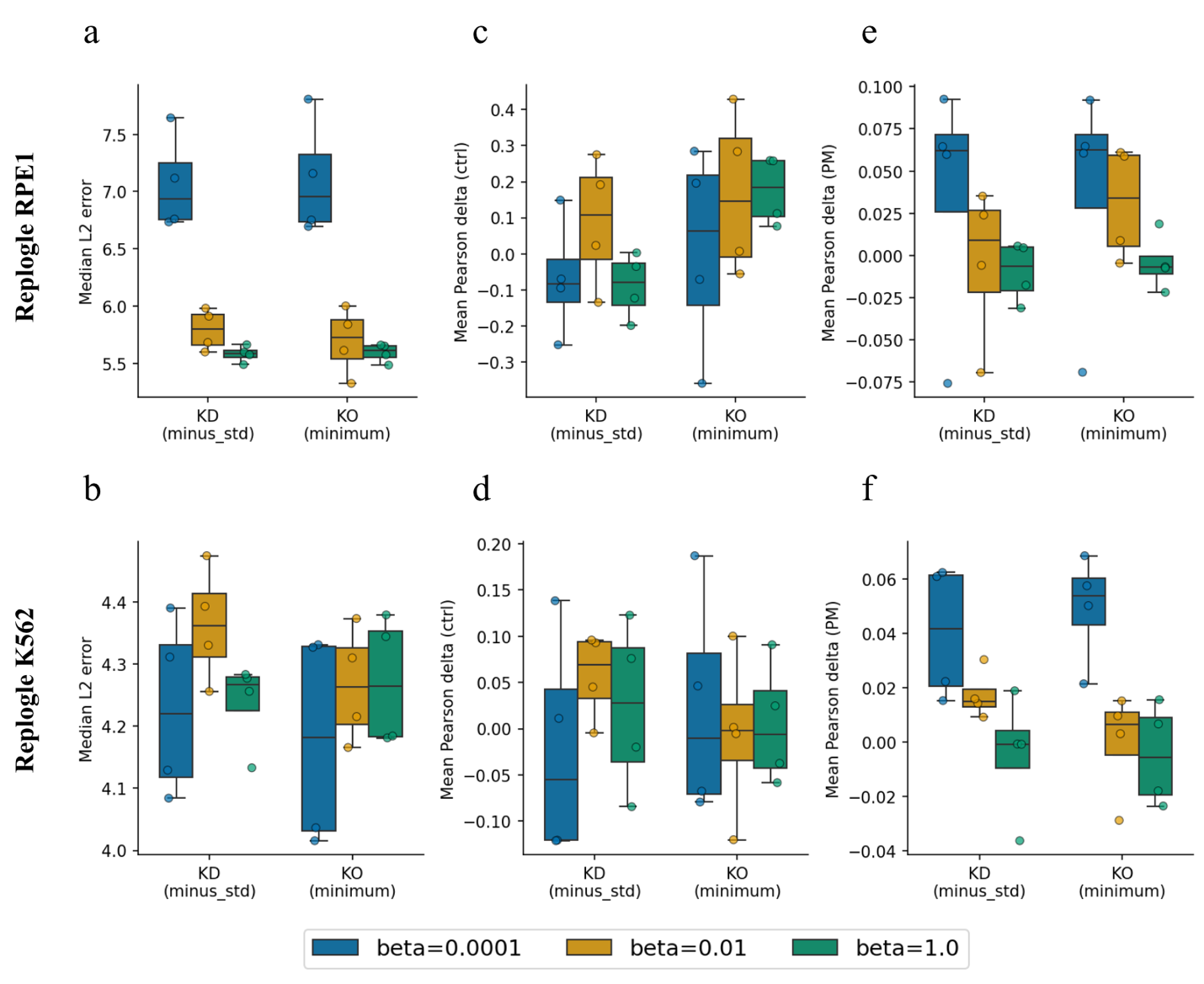

**Supplementary Figure 22. Evaluation of *in silico* input perturbations on single cell CRISPR KD data, larger model (hid=512, lat = 128). a,b.** Median L2 test error across 4 folds under different regularization schemes. **c, d.** Mean Pearson delta measured as the correlation between predictions and observations after subtracting the expression in the control condition. **e, f.** Mean Pearson delta measured as the correlation between predictions and observations after subtracting the mean expression profile across perturbations. This figure shows that a larger model architecture did not yield better predictions, i.e. model limitations were not driven by the reduced model size.

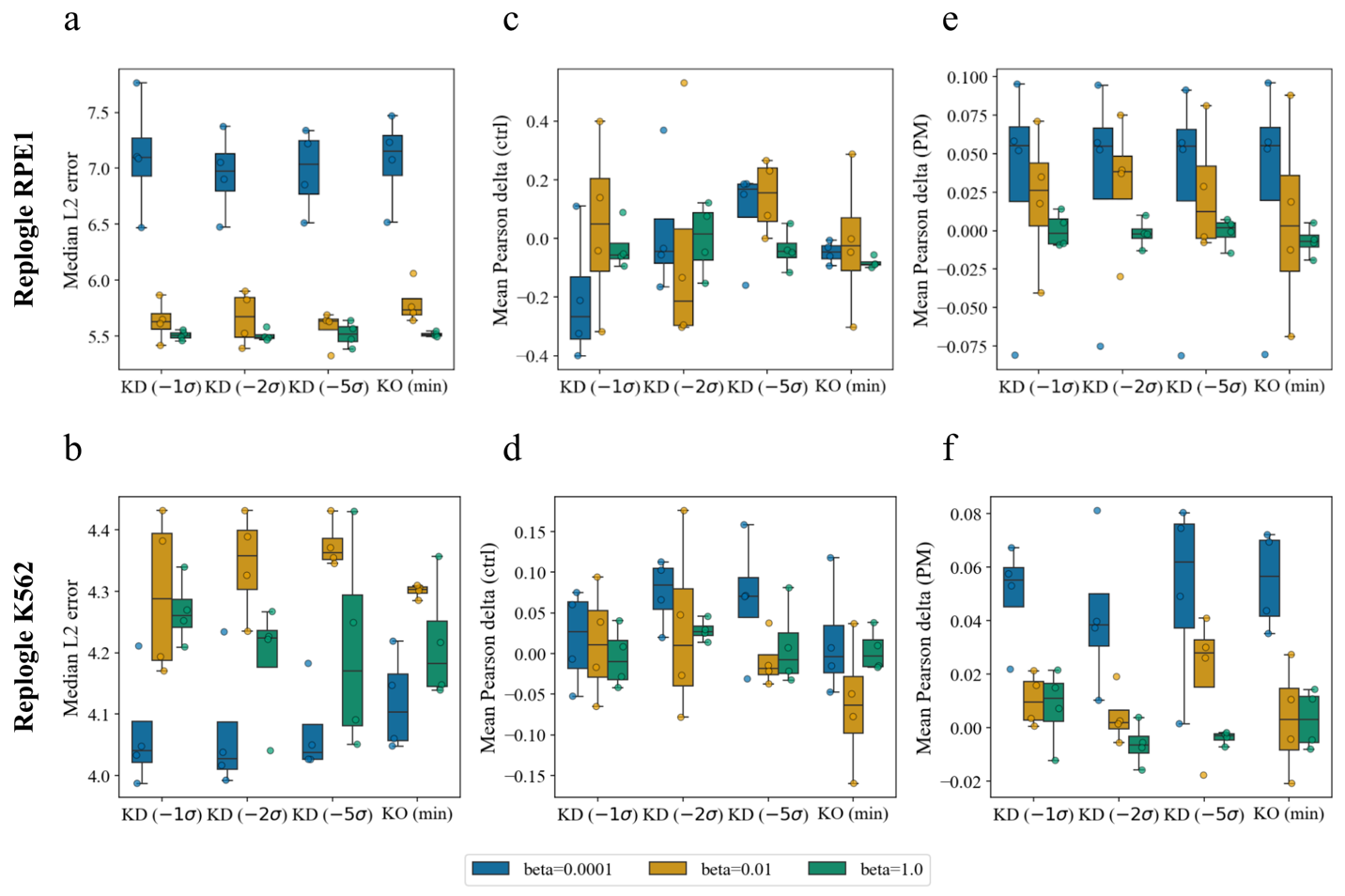

**Supplementary Figure 23. Sensitivity analysis of predicted changes across perturbation magnitudes. a,b.** Median L2 test error across 4 folds under different perturbation magnitudes. **c, d.** Mean Pearson delta measured as the correlation between predictions and observations after subtracting the expression in the control condition. **e, f.** Mean Pearson delta measured as the correlation between predictions and observations after subtracting the mean expression profile across perturbations. The impact of the perturbation magnitude in the predictions is less pronounced than that of the regularization regime.

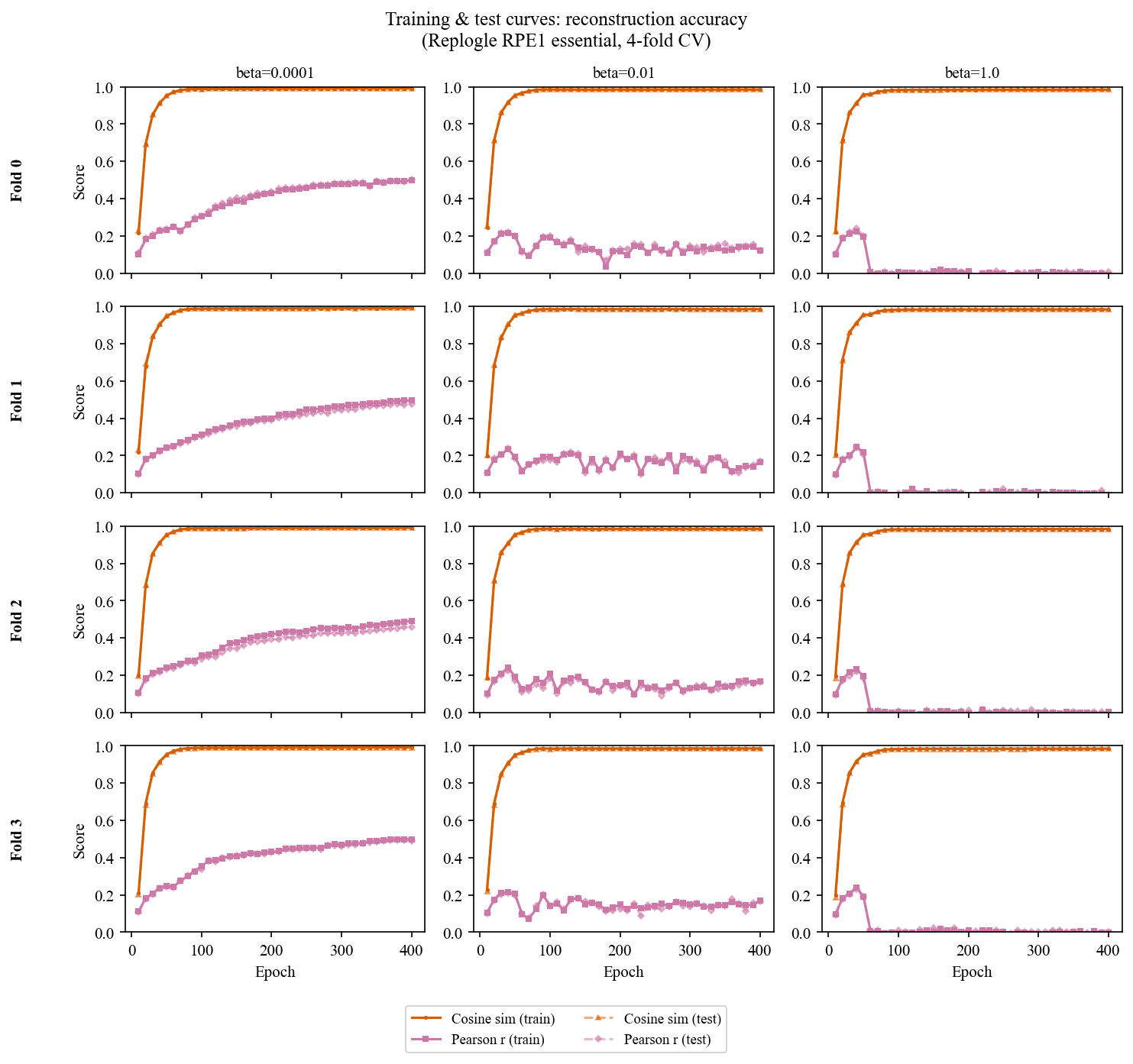

**Supplementary Figure 24. Evolution of the reconstruction quality across folds for Replogle’s RPE1 dataset.** The orange curve tracks the cosine similarity between input and reconstructed profiles. The pink curve tracks mean Pearson correlation between reconstructed and input expression levels, gene-wise. Both metrics are computed on the evaluation set of genes, i.e. the 1000 genes with the highest expression in control cells. Model architecture was the same as for IBDMDB data, with 64 hidden nodes and 16 latent nodes. We can observe the posterior collapse at the high regularization regime. The cosine similarity gets close to one while individual gene Pearson correlations drop to zero, as the model’s incentives lead it to predict the mean profile across perturbations.

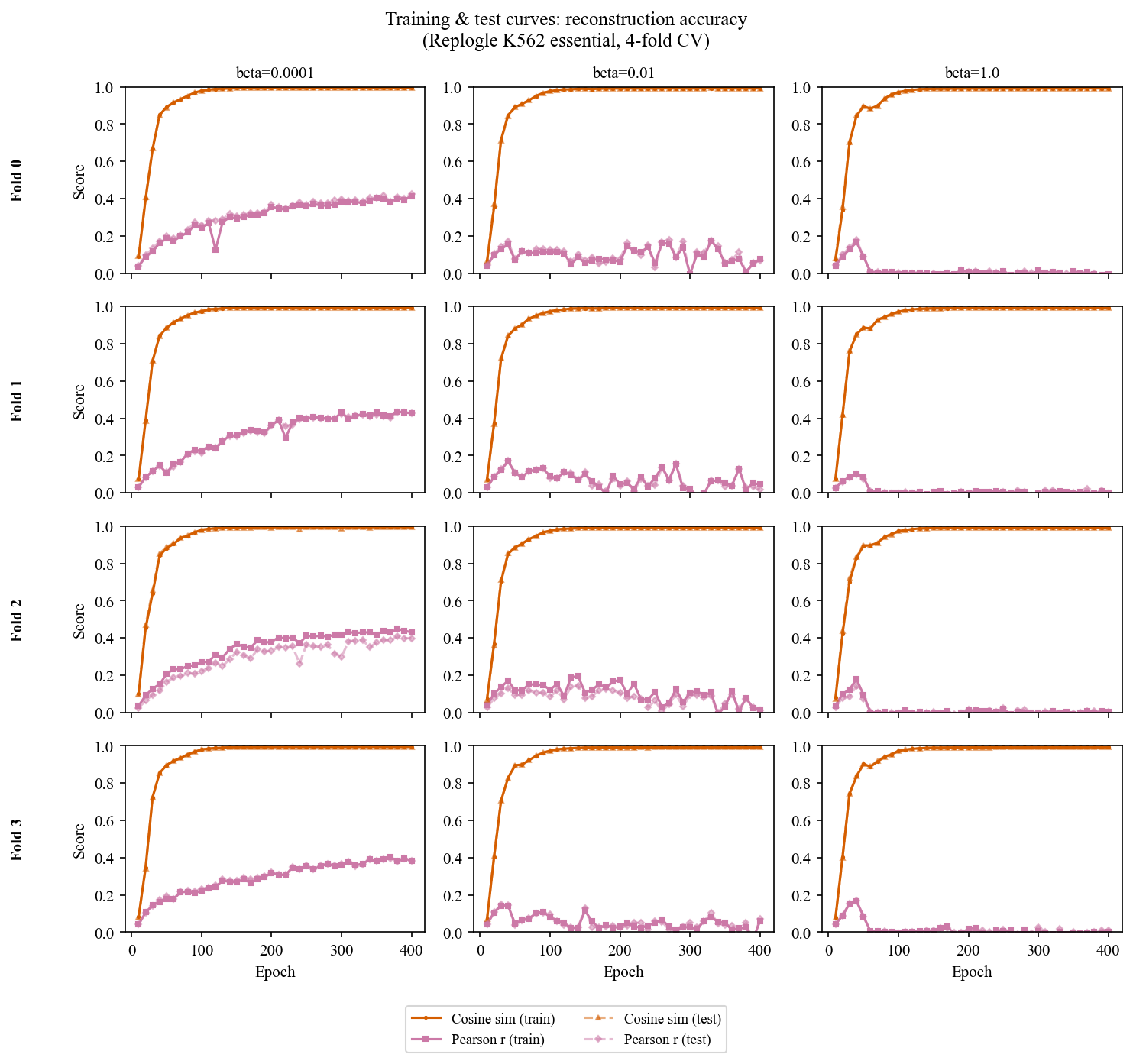

**Supplementary Figure 25. Evolution of the reconstruction quality across folds for Replogle’s K562 dataset.** The orange curve tracks the cosine similarity between input and reconstructed profiles. Mean Pearson correlation between reconstructed and input expression levels, per gene. Both metrics are computed on the evaluation set of genes, i.e. the 1000 genes with the highest expression in control cells. Model architecture was the same as for IBDMDB data, with hid=64 and lat=16.

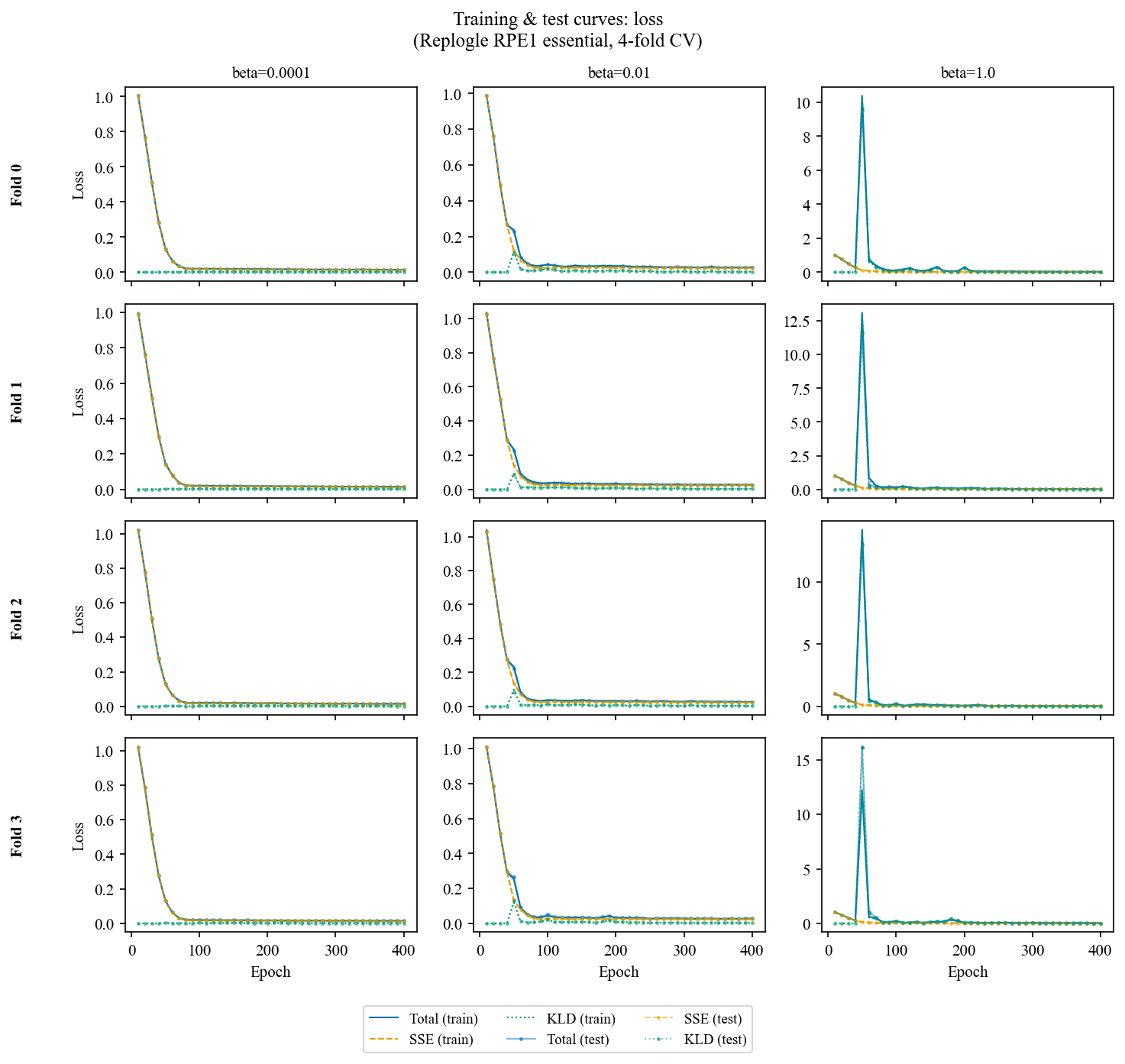

**Supplementary Figure 26. Evolution of the losses across folds for Replogle’s RPE1 dataset.** The same KLD warm-up was applied as for IBDMDB analyses, i.e. introducing the regularization term in fractions at epochs 50, 100 and 150.

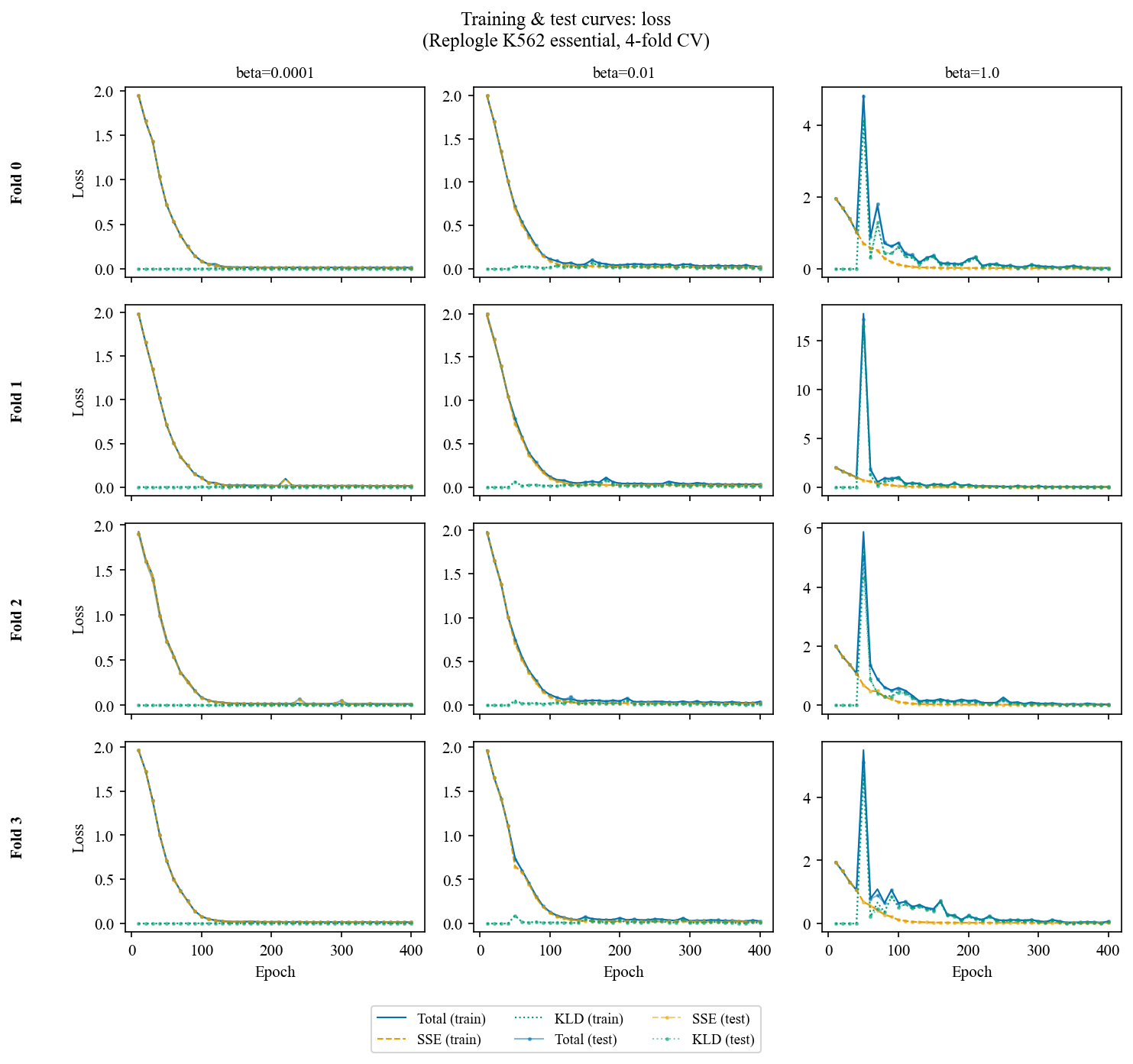

**Supplementary Figure 27. Evolution of the losses across folds for Replogle’s K562 dataset.** The same KLD warm-up was applied as for IBDMDB analyses, i.e. introducing the regularization term in fractions at epochs 50, 100 and 150. This is really pronounced at high beta for when it is first introduced (loss peak).
